# Structural Insights into Curli CsgA Cross-β Fibril Architecture Inspired Repurposing of Anti-amyloid Compounds as Anti-biofilm Agents

**DOI:** 10.1101/493668

**Authors:** Sergei Perov, Ofir Lidor, Nir Salinas, Nimrod Golan, Einav Tayeb-Fligelman, Maya Deshmukh, Dieter Willbold, Meytal Landau

## Abstract

Curli amyloid fibrils secreted by *Enterobacteriaceae* mediate host cell adhesion and contribute to biofilm formation, thereby promoting bacterial resistance to environmental stressors. Here, we present crystal structures of amyloid-forming segments from the major curlin subunit, CsgA, revealing steric zipper fibrils of tightly mated β-sheets, demonstrating a structural link between curli and human pathological amyloids. We propose that these cross-β segments structure the highly robust curli amyloid core. D-enantiomeric peptides, originally developed to interfere with Alzheimer’s disease-associated Amyloid-β, inhibited CsgA fibrillation and reduced biofilm formation in *Salmonella typhimurium*. Moreover, CsgA fibrils cross-seeded fibrillation of Amyloid-β, providing further support for the proposed structural resemblance and potential for cross-species amyloid interactions. In this study, we provide structural insights into curli formation, offer a novel strategy for disrupting amyloid-structured biofilms, and hypothesize on the formation of self-propagating prion-like species originating from a microbial source that could influence neurodegenerative diseases.

**Significance:** Atomic resolution structural insights into the biofilm-associated curli amyloid fibril secreted by *Enterobacteriaceae* revealed elements of fibrillar architecture conserved between bacterial and human amyloids. This inspired us to repurpose anti-amyloid drugs designed to target human pathological amyloids as a novel class of anti-biofilm agents. Moreover, the results provide a molecular basis for understanding interspecies cross-seeding of amyloids through the generation of prion-like agents by molecular mimicry. This raises concerns regarding human exposure to exogenous sources of amyloids, such as contaminated food and amyloid-secreting microbes. Overall, we provide a novel framework for investigating interspecies amyloid interactions at the molecular level and offer novel insights into mechanisms which may underlie the evolutionary and etiological relationships between the human and microbial amylomes.

## Introduction

Amyloid formation has traditionally been viewed as a hallmark of protein misfolding diseases (1). In diseases such as Alzheimer’s and Parkinson’s, amyloids form highly stable cross-β fibrils of mated β-sheets with β-strands situated perpendicular to the fibril axis (2-21). In a groundbreaking study, Chapman and coworkers discovered that *Escherichia coli* secrete extracellular fibers called curli that biochemically and biophysically resemble human pathological amyloidsm (22). Curli fibrils mediate host cell adhesion and invasion, lead to immune system activation, and scaffold protective bacterial communities known as biofilms (23-26). In contrast to disease-associated amyloids, curli and other functional amyloids are produced via dedicated and tightly regulated cellular processes (27-29). Functional amyloids have been identified in all kingdoms of life, demonstrating a ubiquitous role for amyloids in physiological processes (22, 23, 26, 30-33). In microbes, amyloids often serve as virulence determinants involved in aggressive infections and are thus attractive drug targets (26, 34).

In contrast to the study of eukaryotic amyloids, the study of bacterial amyloids has proceeded in a vacuum of high-resolution structural knowledge for years (2-21). We previously determined the first crystal structure of a bacterial amyloid fibril, the phenol soluble modulin α3 (PSMα3) peptide secreted by the pathogenic *Staphylococcus aureus* (35). The structure revealed a fundamental polymorphism of the amyloid fold, showing that α-helices can replace the β-strands stacked perpendicular to the fibril axis, forming cross-α fibrils which are toxic to human cells (35). In contrast, a truncated PSMα3 showed antibacterial activity and revealed two extremely polymorphic and atypical β-rich fibril architectures, a β-rich hexameric arrangement with channels along the fibril axis and an out-of-register β-sheet arrangement (36). Two biofilm-structuring members of the PSM family, PSMα1 and PSMα4, form canonical cross-β amyloid fibrils with β-sheets tightly mated through steric zipper interfaces (36) that likely contributes to the stability of the biofilm matrix (37). Overall, current information on bacterial functional amyloids indicates that structural plasticity underlies functional diversity (35, 36, 38). In this study, we provide structural insights into curli fibril architecture.

The two key players in curli fibril formation are CsgA and CsgB, the major and minor curlin subunits, respectively (29). CsgB nucleates CsgA fibrillation in-vivo via interactions with soluble and unstructured CsgA monomers secreted to the outer bacterial membrane (27-29). This specific nucleation process likely ensures fibril homogeneity and integrity (39). Amyloid fibrils composed of CsgA are further resistant to chemical and proteolytic degradation (22, 40-42). CsgA, a 151-residue long protein, consists of five imperfect sequence repeats (R1-R5), defined by regularly spaced serine (Ser), glutamine (Gln) and asparagine (Asn) residues. The first and the last repeats (R1 and R5) form amyloid fibrils in isolation and are critical to CsgA seeding and nucleation by CsgB(43), while the other repeats (R2-R4) contain ‘gatekeeper’ residues that temper the amyloidogenicity of CsgA (41, 44, 45). CsgA fibrils bind the amyloid indicator dyes thioflavin T (ThT) (46) and congo red (CR) and can be visualized surrounding the bacteria via transmission electron microscopy (TEM) (22, 45, 47). The X-ray fibril diffraction pattern of CsgA shows reflections at 4.6-4.7 Å and 8-10 Å, indicating a structural spacing typical of cross-β architecture (42, 47) reminiscent of human pathological amyloids. Solid-state NMR data has previously suggested that recombinant CsgA adopts the same structure as native curli isolated from the bacteria, validating much of the biophysical information obtained from recombinant CsgA (48).

Molecular structures of curli-associated proteins are needed to better understand how curli biogenesis is controlled. Structures are available for few curli accessory components including solution NMR structures of the periplasmic accessory protein CsgE (49) and the adaptor protein CsgF (50), as well as crystal structures of the periplasmic chaperone CsgC (51), which inhibits intracellular CsgA fibrillation (28, 52), and the outer membrane secretion channel CsgG (53, 54). Despite attempts to obtain structural information on curli fibrils, atomic resolution structures of CsgA and CsgB have remained elusive due to limitations of some structural methods in contending with the intrinsic properties of full-length amyloid fibrils, which are insoluble and often polymorphic and partially disordered in nature (6, 14, 21, 48). We thus adopted the reductionist approach of looking for amyloid spine segments which are suggested to form the structured backbone of the fibril (21).

We obtained four atomic structures of amyloidogenic six-residue segments from CsgA, three of which we propose may serve as amyloid spines. These spines contain residues essential for fibrillation and adopt elongated and unbranched fibrils that bind ThT, the amyloid-binding dye (46). The structures of these three spines are reminiscent of those of human pathological amyloids, forming tightly paired β-sheets that belong to class 1 steric-zippers, classified according to the organization of the β-strands and β-sheets (12, 21). The structural similarity between fibrillar spine segments derived from CsgA and those derived from human pathological amyloids prompted us to investigate whether fibrillation inhibitors designed against human amyloids could also inhibit curli formation. Accordingly, we found that two D-enantiomeric peptides, originally designed to interfere with the formation of oligomers of Alzheimer’s disease-associated Aβ (55-65), inhibited the fibrillation of CsgA spines as well as of full-length CsgA and reduced biofilm formation in curli-expressing *Salmonella typhimurium* in dose-dependent manners. These results provide structural insights into a biofilm-related amyloid and pave the way for the rational development of anti-microbial drugs targeting amyloid-structured biofilms.

## Results

### Atomic resolution structures of CsgA spine segments reveal canonical steric-zippers characteristic of pathological amyloids

To investigate structural features of CsgA, we identified potential amyloid-forming segments that may function as structured spines of CsgA fibrils. These segments were selected by combining computational data predicting regions of amyloidogenic propensity (66-70). We focused on _45_LNIYQY_50_ and _47_IYQYGG_52_ from the R1 repeat, _137_VTQVGF_142_ from the R5 repeat, and _129_TASNSS_134_ from the R4-R5 loop (sequence positions are indicated as subscript based on the sequence of CsgA with a Uniprot accession number P28307). _129_TASNSS_134_ was selected as a control sequence as it was computationally predicted to be amyloidogenic but is located in a region not implicated in fibrillation (43). TEM micrographs demonstrated that all four segments formed fibrils (Fig. S1) in accordance with the predictions of their amyloidogenic propensities. However, while _45_LNIYQY_50_ (R1), _47_IYQYGG_52_ (R1), and _137_VTQVGF_142_ (R5) formed amyloid-like unbranched and elongated fibrils, the _129_TASNSS_134_ (R4-R5 loop) segment formed wide and atypical structures, far removed from amyloid fibrils (Fig. S1). The _45_LNIYQY_50_ (R1), _47_IYQYGG_52_ (R1), and _137_VTQVGF_142_ (R5) segments bound the amyloid indicator dye ThT, demonstrating dose-dependent amyloid fibrillation curves with short lag times (Fig. S2). _45_LNIYQY_50_ polymerized with the shortest lag time. The _129_TASNSS_134_ segment from the R4-R5 loop did not bind ThT (Fig. S2).

To obtain atomic-resolution structural insight into curli fibrils we solved the crystal structures of the four segments (Fig. 1, Figs. S3-S6, Table S1). Notably, _45_LNIYQY_50_ and _129_TASNSS_134_ formed crystals that diffracted well only when mixed with the _134_TAIVVQ_139_ segment from the R5 repeat of the nucleator protein CsgB prior to crystallization. Namely, TAIVVQ was not present in the crystals, only aided in crystallization. It is possible that this phenomenon is relevant to the mechanism of heteronucleation of CsgA by CsgB in-vivo (29). _45_LNIYQY_50_ and _47_IYQYGG_52_ are overlapping segments from the R1 repeat and adopt very similar structures. Both segments, as well as _137_VTQVGF_142_ (R5), adopt classical amyloid steric-zipper architecture with two possible dry interfaces between paired β-sheets (Fig. 1). The β-strands are oriented parallel to each other along the β-sheets. These three structures are class 1 steric zippers as defined by Sawaya and Eisenberg according to the organization of the β-strands and β-sheets (12, 21). In each of these dry interfaces, the chemical properties governing fibril stability, i.e., buried surface area and shape complementarity between sheets, resembles those of eukaryotic steric-zipper structures (Table S2). Correspondingly, the three segments formed fibrils that bound the amyloid-indicator dye ThT (Figs. S1-S2). In agreement with these observations, previous alanine scanning mutagenesis analyses showed that Gln49, Asn54, Gln139, and Asn144 in CsgA are essential for curli assembly (43). _45_LNIY**Q**Y_50_, _47_IY**Q**YGG_52_, and _137_VT**Q**VGF_142_ each contained one of these essential residues (marked in bold).

**Figure 1.**
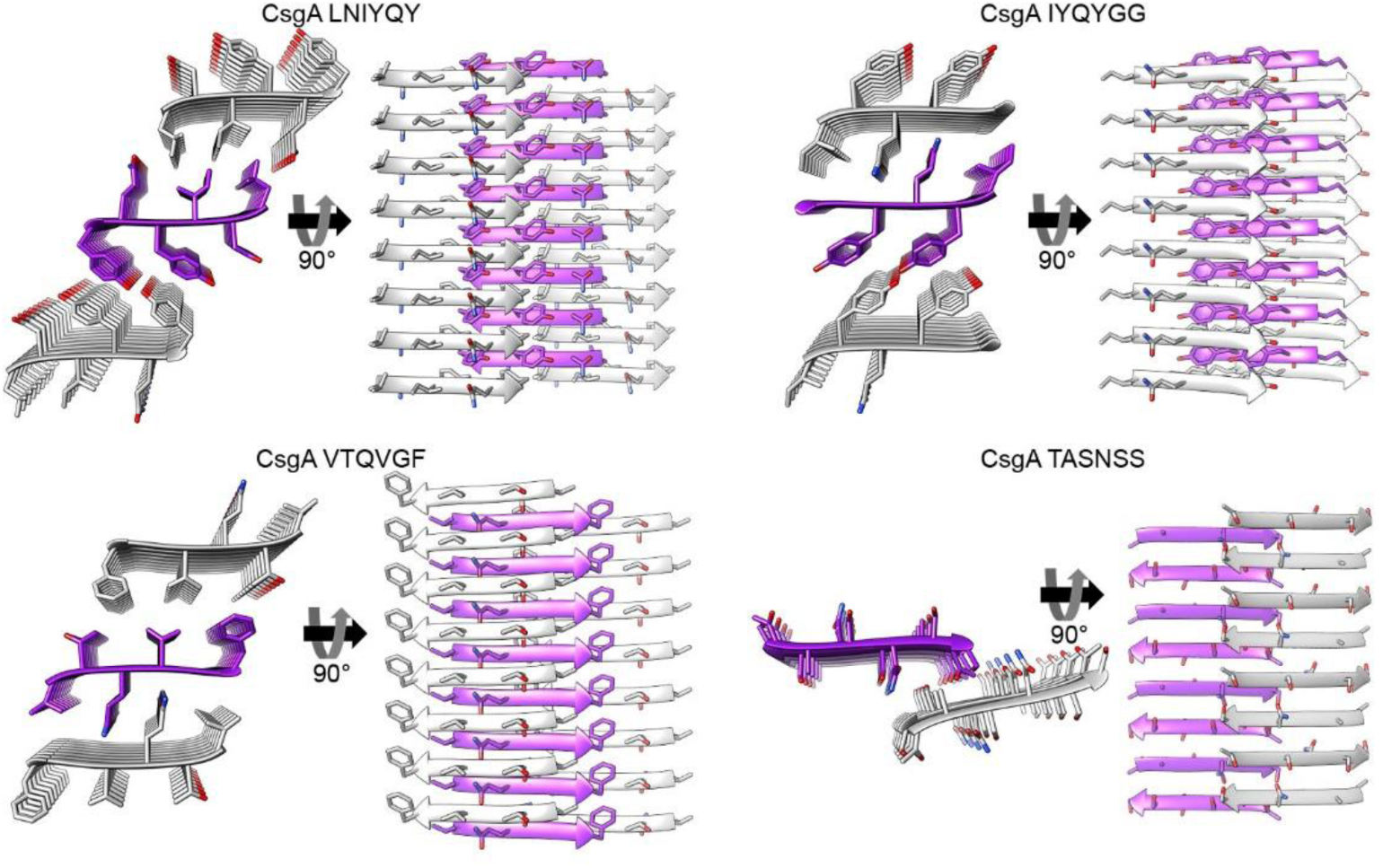
Crystal structures of CsgA spine segments. High-resolution crystal structures of CsgA segments are shown. In each structure, the left view is down the fibril axis with residues shown as sticks, and the right view is perpendicular to the fibril axis, with β-strands, shown as ribbons, run horizontally. Eight layers of β-strands are shown while actual fibrils contain thousands of layers. The carbons of each β-sheet are colored either gray or purple; heteroatoms are colored by atom type (nitrogen in blue, oxygen in red). The _45_LNIYQY_50_, _47_IYQYGG_52_ and _137_VTQVGF_142_ segments formed the classical class 1 steric zipper architecture of tightly mated parallel β-sheets with individual subunits (peptides) situated perpendicular to the fibril axis. Two possible dry interfaces in the crystal packing are shown. The _129_TASNSS_134_ segment formed extended β-strands that stack in an anti-parallel manner and most closely resemble class 8 steric zippers but with a small contact area between two facing β-sheets and lacking a dry interface. Figs. S3-S6 provide a detailed description of the four crystal structures.

_129_TASNSS_134_, the segment derived from the R4-R5 loop, was selected as a control because it was computationally predicted to form amyloid but is not known to participate in CsgA fibrillation. In contrast to the three other peptides, _129_TASNSS_134_ does not contain any residues that have been shown to act as sequence determinants of CsgA fibrillation (43). Moreover, Ser133 is nonessential for fibrillation (43). Unlike the three spine segments that adopt tightly packed steric zipper architectures, _129_TASNSS_134_ adopts anti-parallel β-sheets lacking a dry interface between mated sheets. The packing of the β-sheets most closely resembles a class 8 steric zipper (12), with a truncated interface between the two facing β-sheets. In agreement with the atypical crystal structure of _129_TASNSS_134_, it formed ribbon-like structures, which are exceptionally flat and wide compared to canonical amyloid fibrils (Fig. S1), and did not bind ThT (Fig. S2). This demonstrates that not all segments that are predicted by servers to have high aggregation propensities form steric zippers. Here, only those segments which are likely to play a role in nucleating fibrillation according to biochemical evidence formed steric zippers.

### CsgA fibrils share structural characteristics with cross-β amyloids and seed the fibrillation of human Aβ

The structural features of full-length CsgA fibrils were analyzed using attenuated total-internal reflection Fourier transform infrared (ATR-FTIR) spectroscopy and showed a main peak at 1617 cm^−1^ corresponding to a rigid cross-β amyloid architecture (71-73) (Fig. S7). Furthermore, in accordance with a previous report (74), CsgA seeds are able to accelerate the fibrillation of Alzheimer’s disease-associated Aβ_1-40_ and Aβ_1-42_, which have been shown to adopt canonical cross-β fibril architectures (8) (Fig. S8). Epitaxial heteronucleation necessitates some degree of structural similarity between participating amyloids, suggesting that full-length CsgA adopts a fibril architecture sufficiently similar to that of Aβ_1-40_ and Aβ_1-42_ to effectively template fibril elongation (74, 75). Nucleation is expected to be specific, as for example, islet amyloid polypeptide protein fibrils were unable to seed CsgA (76).

### CsgA shares fibrillation inhibitors with Aβ

We next sought to investigate whether known inhibitors of human pathological amyloid formation would effectively inhibit CsgA fibrillation on the basis of this proposed structural similarity. We tested a group of synthetic D-enantiomeric peptides (referred to here as D-peptides) designed against Aβ (55-65). Two of the D-peptides tested, ANK6 and DB3DB3 (60), were found to inhibit CsgA fibrillation in dose-dependent manners. Specifically, freshly purified recombinant CsgA show the characteristic ThT amyloid-fibrillation kinetics curve with a very short lag time followed by rapid aggregation, while the addition of the D-peptides resulted in lower fluorescence signal and a longer lag-time, indicative of delayed fibril formation (Fig. 2A and Fig. S9). At 1:5 molar ratios of CsgA to each inhibitor, ANK6 induces a longer lag time in CsgA fibrillation than did DB3DB3, but they eventually reach similar maximal fluorescence intensities, which are significantly lower than that of CsgA without the D-peptides (Fig. 2A). This data may reflect the ability of the D-peptides to induce different morphologies of the CsgA fibrils, or interfere with one or more microscopic processes underlying the kinetics of CsgA fibrillation. Dose-dependent inhibition of CsgA fibrillation by the D-peptides is shown in Fig. S9, demonstrating that a significant inhibitory effect is observed already at 1:1 molar ratios of CsgA to the D-peptides. The TEM micrographs correspondingly showed mostly amorphous aggregates or co-precipitates of CsgA in the presence of ANK6 and DB3DB3 compared to fibrils of CsgA alone (Fig. 2C). Both ANK6 and DB3DB3 were more potent fibrillation inhibitors than D3 (Fig. 2A and Fig. S9), a prototype D-peptide previously shown to reduce amyloid deposits and inflammation and improve cognition in transgenic mouse models for Alzheimer’s disease (56). In agreement with the hypothesis that the steric-zipper segments serve as spines of CsgA fibrils, we found that the D-peptides also inhibited the fibrillation of the R1 and R5 spine segments in dose-dependent manners (Fig. S10). Moreover, the same inhibitory series for the D-peptides (ANK6>DB3DB3>D3) was observed for the spines. In contrast to CsgA and its spine segments, ANK6 and DB3DB3 did not affect the fibrillation of the PSMα3 peptide secreted by the pathogenic *S. aureus* bacterium, which forms cross-α amyloid-like fibrils (35) (Fig. 2B), suggesting that inhibition is dependent on the secondary structure of the fibril.

**Fig. 2.**
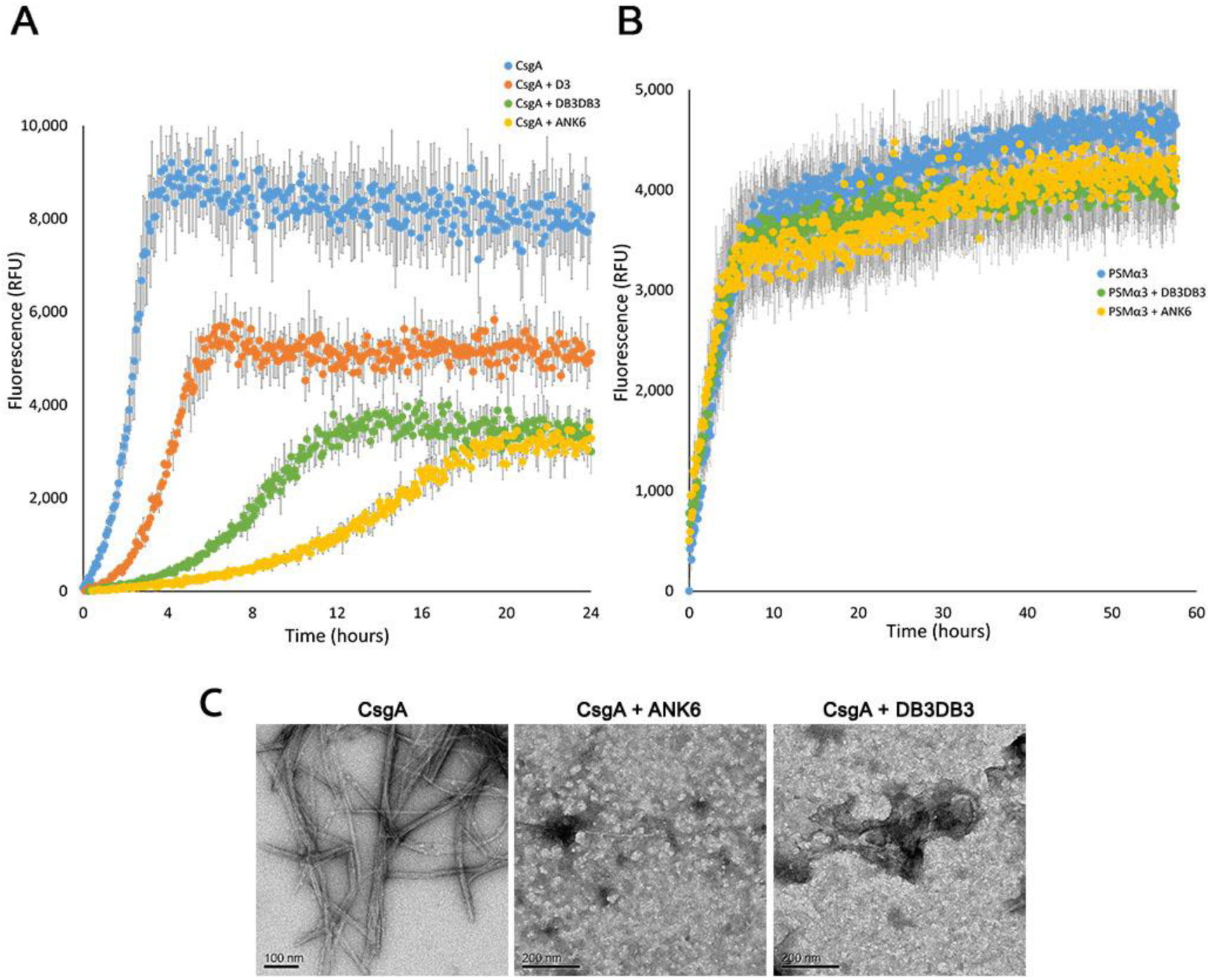
D-peptides inhibit fibrillation of CsgA but not of PSMα3. The graphs show mean fluorescence readings of triplicate ThT measurements of CsgA (**A**) or PSMα3 (**B**) with or without the D-peptides at a 1:5 molar ratio, respectively. Error bars represent standard error of the mean calculated from triplicates. ANK6 and DB3DB3 delay fibril formation of CsgA and reduce the fluorescence signal, while demonstrating no effect on PSMα3 fibrillation. D3 shows a minor effect on CsgA (**A**). (**C**) Electron micrographs of CsgA incubated overnight showing the formation of fibrils. In the presence of ANK6 or DB3DB3 (1:10 molar ratio of CsgA to D-peptides), only amorphous aggregates are observed. Scale bars are indicated for each micrograph.

The effect of ANK6 and DB3DB3 on the secondary-structure transition of CsgA during fibrillation was assessed using time-dependent circular dichroism (CD) spectroscopy (Fig. S11). The CD spectra of freshly purified recombinant CsgA displays a typical random coil configuration with a detected minimum around 195 nm. After six hours of incubation, the spectra of CsgA shows a transition to a well-ordered β-sheet structure with a distinctive maximum near 198 nm and minimum near 218 nm. The timescale of this transition is similar to that observed in previous work (47). The CD spectra of CsgA incubated with ANK6 (at 1:5 molar ratio) displayed that CsgA retained random coil configuration along 18 hours of incubation (Fig. S11), indicating inhibition of CsgA fibrillation.

CsgA fibrils are resistant to sodium dodecyl sulfate (SDS) solubilization (47, 77). We therefore examined the effect of the D-peptide inhibitors on the SDS soluble-to-insoluble shift of CsgA by assessing its migration with SDS polyacrylamide gel electrophoresis (SDS-PAGE) (Fig. 3 and Fig. S12). We first assessed the oligomeric state of freshly purified recombinant CsgA using size exclusion chromatography coupled with multi angle light scattering (SEC-MALS) (Fig. S13). This analysis showed that the majority of freshly purified CsgA is in the monomeric state. A minor population exists as hexameric oligomers. The presence of some oligomers at these early time points would not be surprising considering the rapid fibrillation kinetics observed for CsgA, but it is still not clear whether these hexamers play a role in CsgA fibrillation process. Interestingly, the formation of pentamers or hexamers as the smallest populated assembly species has been described for Aβ as well (78). Freshly purified recombinant CsgA incubated for a few hours forms SDS-resistant fibrils that do not migrate in SDS-PAGE, in contrast to the freshly purified soluble CsgA (Fig. 3 and Fig. S12). The effect of the D-peptide inhibitors on the fibrillation of CsgA (at 5:1 molar ratio, respectively) is evident: DB3DB3, and to a greater extent ANK6, stabilized the soluble CsgA state. This suggests a rapid and robust effect of the D-peptide inhibitors on CsgA fibrillation in accordance with the ThT, TEM (Fig. 2 and Fig. S9) and CD results (Fig. S11).

**Fig. 3.**
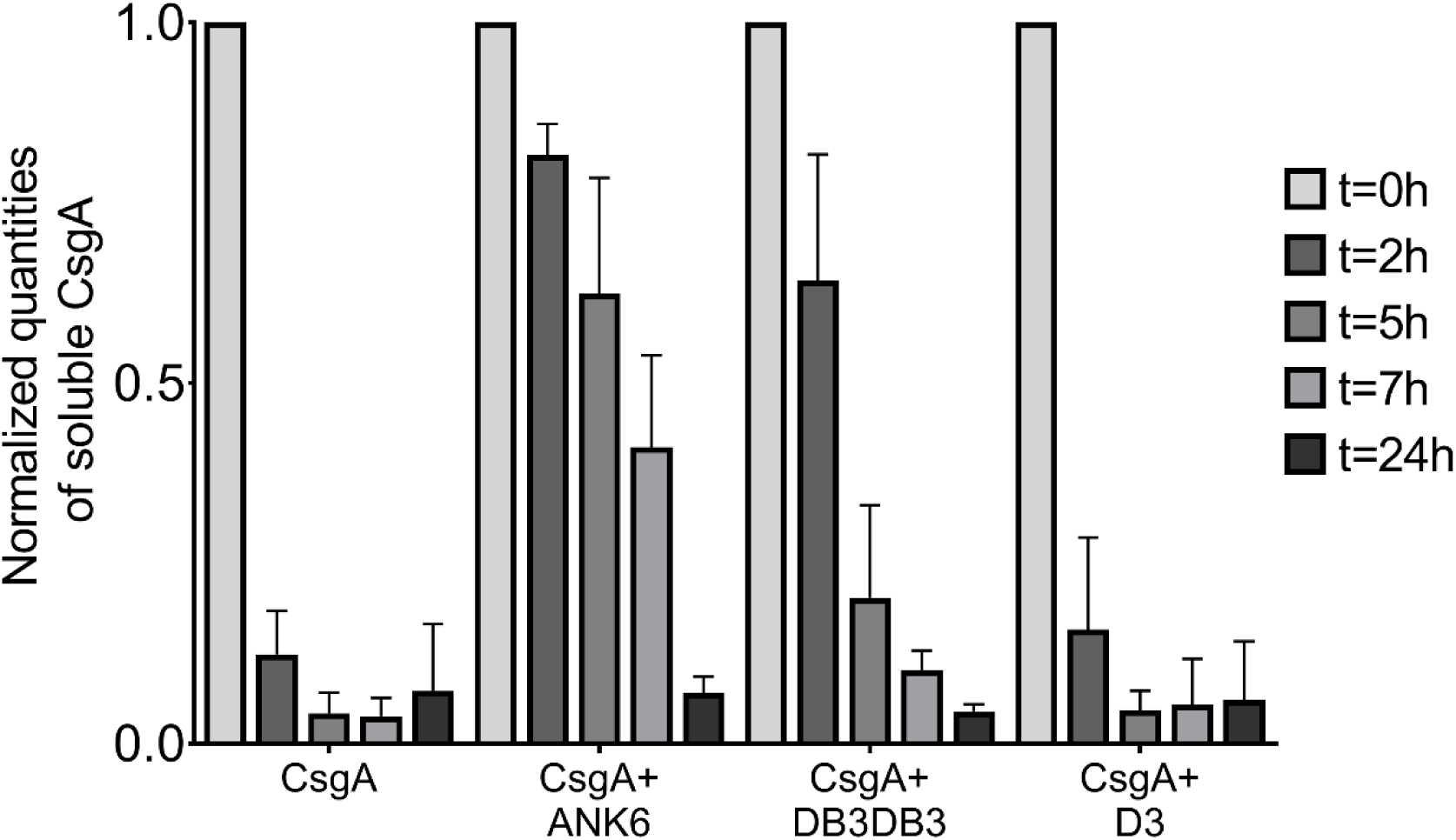
The D-peptides stabilize soluble CsgA and inhibit fibrillation. Quantification of the effect of D-peptide inhibitors on CsgA fibrillation at 5:1 molar ratios based on its ability to migrate as monomer in 15% SDS-PAGE. Each band in the gel (exemplified in Fig. S12) was quantified using a bovine serum albumin (BSA) standard and then normalized to the amount of freshly purified CsgA loaded in each set of migrated bands. CsgA incubated with ANK6 and DB3DB3 showed prolonged soluble state, suggesting inhibition of the rapid formation of SDS-insoluble fibrils. The graph represents three experimental repeats. The experiments were repeated at least three times on different days with similar results.

### The D-peptides reduce biofilm formation of *Salmonella typhimurium*

Treatment with ANK6 and DB3DB3 reduces the total static biofilm biomass of *S. typhimurium* MAE52 strain (a constitutive curli fimbria and cellulose producer) in dose-dependent manners, as shown by reduction in crystal violet staining of the biofilm (Fig. 4). Both DB3DB3 and ANK6 produce significant inhibitory effects at 10µM, with DB3DB3 producing a more pronounced effect. The D3 peptide only affects biofilm formation slightly at 20µM and is less effective than the other two D-peptides, in accordance with the in-vitro results on inhibition of CsgA fibrillation. The dose-dependent effects on the biofilms by these three peptides cannot be attributed to bacteriostatic or bactericidal effects, as no effect on bacterial growth is observed with the addition of up to 150 µM of any of the three peptides (Fig. S14). Confocal microscopy images showing biofilm cells stained with propidium iodide confirm the significant reduction in the formation of an otherwise abundant surface-attached biofilm of *S. typhimurium* by the addition of 10 µM DB3DB3 or ANK6 (Fig. 5). D3 showed no significant effect on the biofilm at 10 µM, in agreement with the crystal violet assay.

**Fig. 4.**
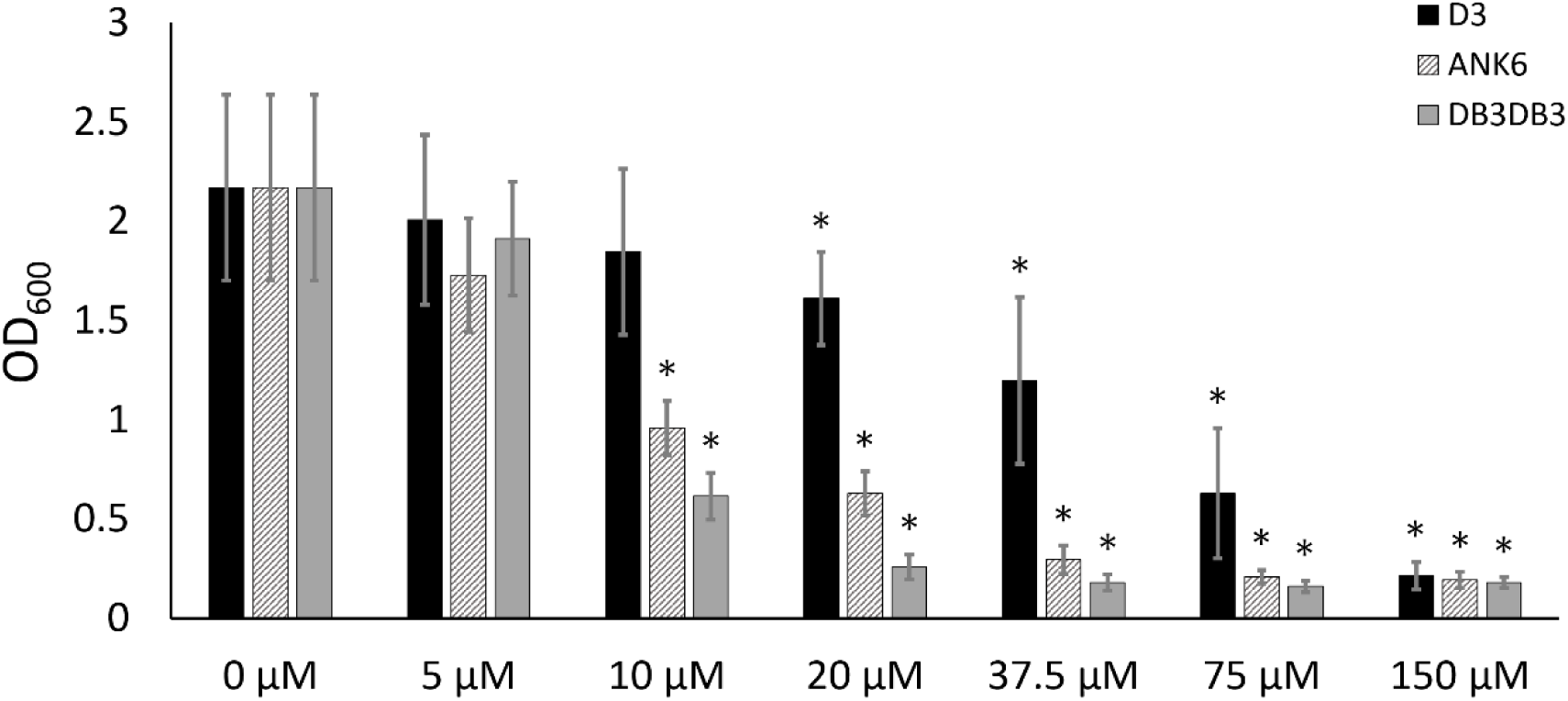
The D-peptides reduce formation of static biofilm of *S. typhimurium*. Quantification of static biofilm production of *S. typhimurium* MAE52 strain without and with the addition of the D-peptides at various concentration using crystal-violet staining measured by optical density at 600 nm (OD_600_). Statistical significance was analyzed by the Mann-Whitney non-parametric test. Standard deviations (S.D) are included (N ≥10). * P<0.05 compared to control (no inhibitor added).

**Fig. 5.**
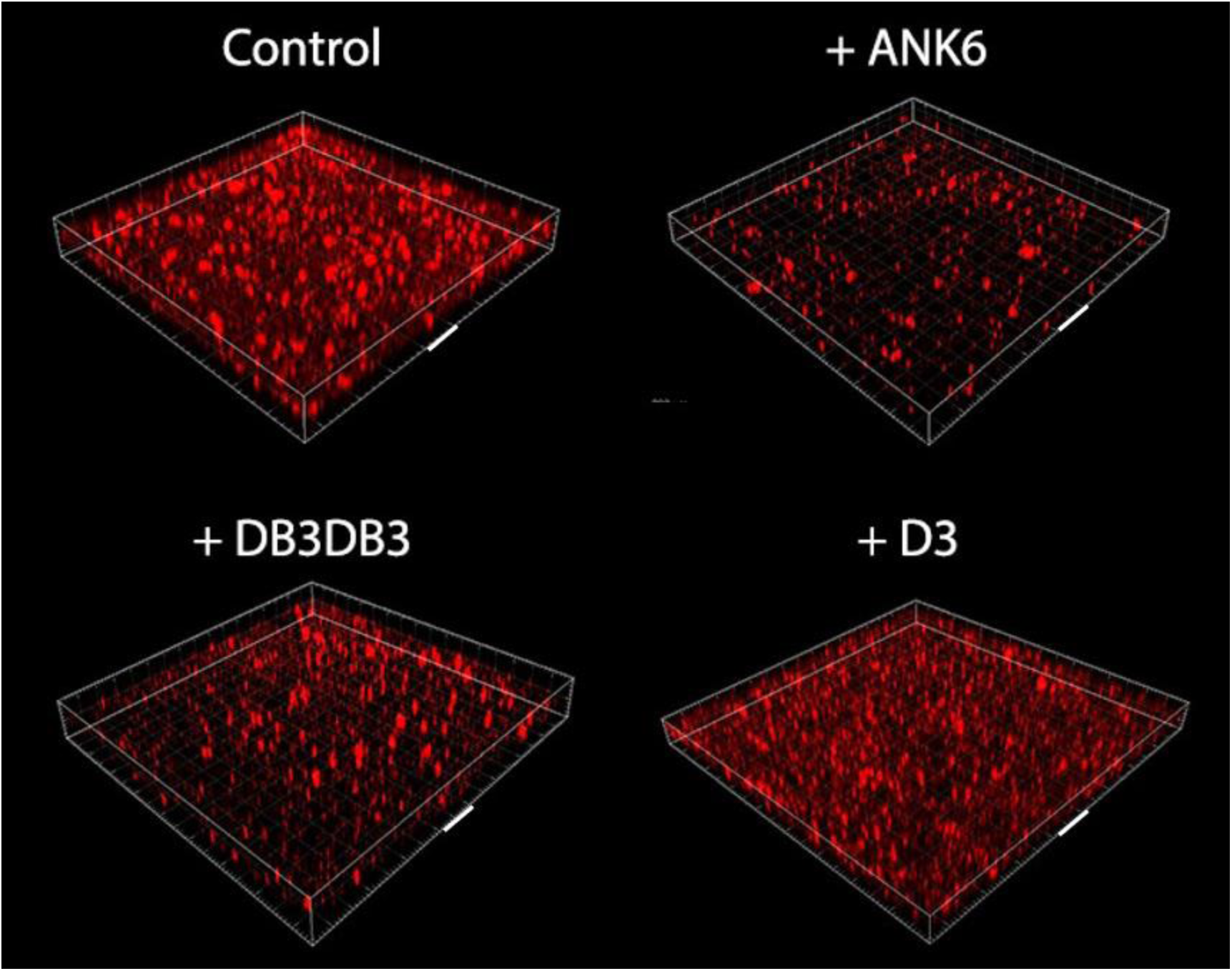
The D-peptides reduce formation of surface-attached static biofilm of *S. typhimurium*. Confocal microscopy images showing surface-attached static biofilm formation using propidium iodide staining without (control) and with the addition of the D-peptides at 10 μM. Each grid box depicts 100×100 µm, the face of one box is marked with a white bar representing 100 µm. These images represent a triplicate of wells, a phenomenon which repeated in at least three independent experiments.

### The D-peptides effects on biofilm formation are amyloid-related

In order to confirm that the effect of the D-peptides on biofilm biomass is related to inhibition of amyloid formation, we used congo red (CR), a dye which is known to stain amyloids, including curliated whole cells (25). Accordingly, growing the *S. typhimurium* MAE52 strain on a CR-supplemented agar resulted in reddish biofilm colonies that adsorbed the dye (Fig. S15A). Addition of ANK6 and DB3DB3 to the bacteria showed a dose-dependent discoloration at the center of the biofilm colony (where initially the suspension was placed on the agar) indicating less CR adsorption (Fig. S15A) and less fibril formation. Since CR also stains secreted cellulose in biofilm (79), we performed further experiments using the *S. typhimurium* MAE150 mutant (MAE52 derived strain) that does not express cellulose in order to verify the effect of the D-peptides on curli fibril formation. In order to comparatively quantify *S. typhimurium* whole-cell curliation with and without the addition of the D-peptides, we removed the colonies from the CR-supplemented agar and measured the concentration of CR that was not adsorbed by the curli-producing bacterial cells but remained in the agar (Fig. 6 and Fig. S15B). DB3DB3 was most effective in reducing curli fibril formation at a dose-dependent manner, followed by ANK6, while D3 only produced an effect at 40 µM. These results are in agreement with the effect of the D-peptides on static biofilm formation (Fig. 4). Overall, ANK6 and DB3DB3 showed inhibitory effect on CsgA fibrillation in-vitro, and on curli production and biofilm formation in the bacteria.

**Fig. 6.**
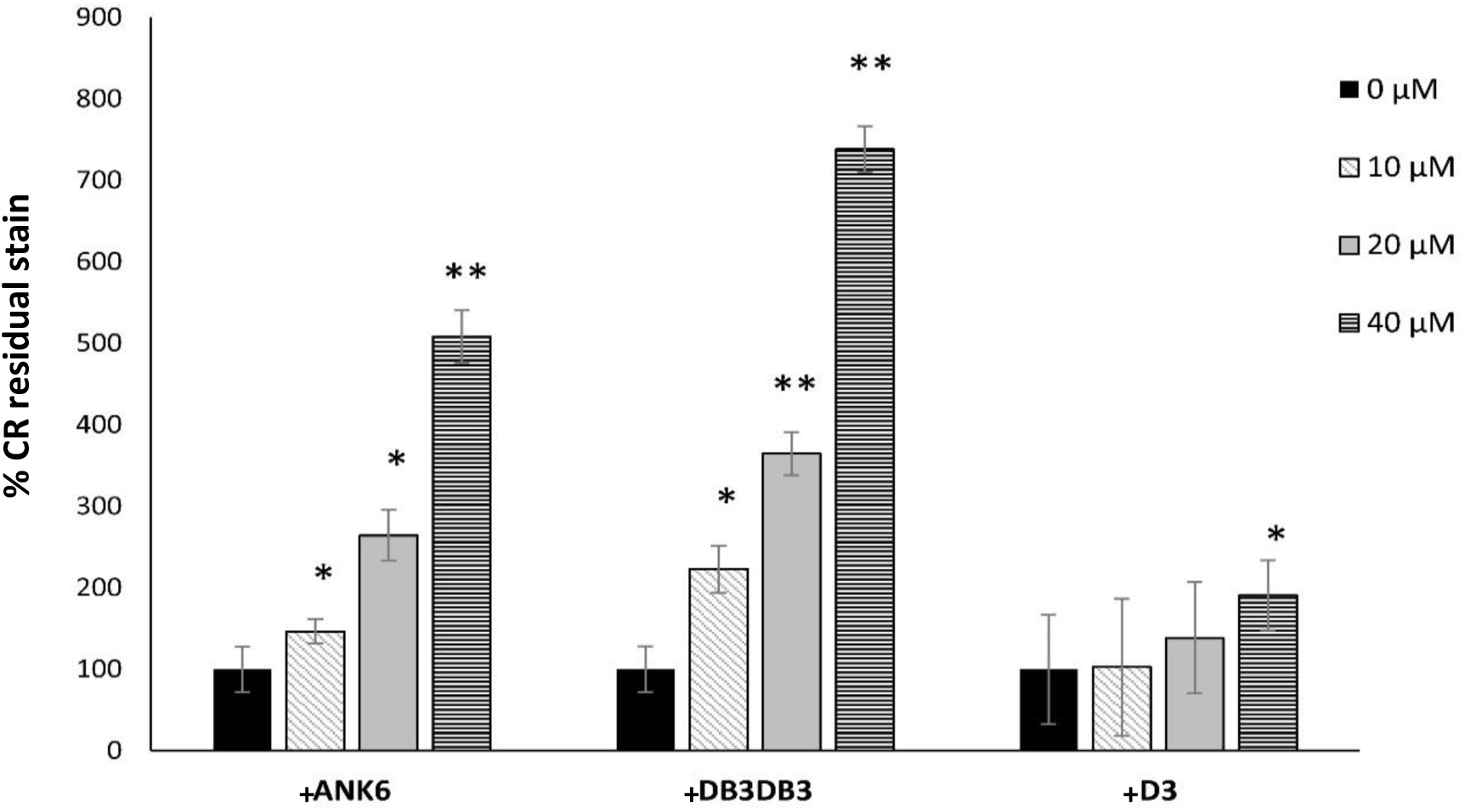
The D-peptides reduce curli production in *S. typhimurium* biofilm. Quantification of the dose-dependent effect of the D-peptides on curli fibril formation in the bacteria. The bars show levels of measured CR residual stain that remained in the agar following the removal of the biofilm colony of the *S. typhimurium* MAE150 cellulose deficient mutant. Control colonies that were not exposed to the D-peptides are assigned 100% CR residual stain that remained in the agar. Higher percentage of CR residual stain indicates less CR adsorption by the colony and hence reduced curli fibril formation. Standard deviations (S.D) are included (N≥12). * P<0.05 and ** P<0.005 compared to control (no inhibitor added). CR plates from one of the experiments are exemplified in Fig. S15B.

## Discussion

### Cross-β steric-zipper fibrils likely stabilize and structure biofilms

The curli biogenesis machinery is designed to secrete, nucleate and elongate extracellular amyloid fibrils that participate in biofilm formation and stability (23-29). While the amyloidogenic properties of the major curlin subunit, CsgA, have been investigated in detail, high-resolution structural information about CsgA fibrils was yet lacking. In this study, we have elucidated atomistic structural features of CsgA spine segments. Specifically, the R1 _45_LNIYQY_50_ and _47_IYQYGG_52_ segments and the R5 _137_VTQVGF_142_ segment formed fibrils that bound the amyloid indicator dye ThT and formed canonical amyloid class 1 steric-zippers (Fig. 1 and Figs. S1-5). These segments contain Gln49 or Gln139 which are critical for fibrillation and cannot even be mutated to asparagine without interfering with curli assembly (43). We postulate that these spine segments scaffold and stabilize the robust curlin amyloid architecture.

The CsgA steric-zipper spine structures closely resemble those of pathological human amyloids (Table S2). Extensive structural studies on pathological amyloids suggest that their cross-β signatures result from steric-zipper-like structures in the fibril spine formed from one or more short segments of the protein (80-82). We therefore expect that the CsgA steric-zipper spine segments comprise the structured core of the CsgA fibril and allow for the formation of cross-β fibrils by full-length CsgA (22, 42, 45, 47). Cross-β ultra-stable structures are known for their extraordinary mechanical properties rendering them as stiff as silk and as strong as steel (83). In bacteria, these fibrils likely stabilize and structure biofilms, thereby rendering the bacterial communities more resilient and resistant to antibiotics (22, 23, 26, 30). Steric zipper structures of spine segments also form the cores of the cross-β fibrils of PSMα1 and PSMα4 found in the biofilm of *S. aureus* (36). Toghther, our findings suggest that the steric zipper may be a structural hallmark of biofilm-associated microbial amyloids as well as their disease-associated counterparts, supporting a linked structural building block from bacteria to human.

### Structural models of full-length CsgA fibrils

A previously proposed structural model of full-length CsgA suggested that the fibrils adopt a β-helix or β-solenoid-like fold. This was based on a model suggested for *Salmonella* CsgA (41) and on information obtained from solid-state NMR and electron microscopy (42, 48) that is more consistent with a β-helix-like structure rather than with in-register parallel β-sheets, although the authors note that the data is insufficient to definitively confirm the adoption of such a structure (42). This arrangement is also supported by a computational model suggesting that in such a β-helical fibril, each turn corresponds to one repeat sequence of CsgA, forming two β-strands connected by a loop creating a “rectangular” hydrophobic core (84). The elongation of the structure is achieved by intermolecular stacking along the fibril axis mediated via the R1 and R5 repeats (84). We herein suggest an alternative model in which fibril formation is nucleated and stabilized via several spine segments in the R1 and R5 repeats. In this model, CsgA adopts a canonical cross-β architecture composed of tightly mated in-register β-sheets, more closely resembling models proposed for human pathological amyloids (8). The lack of polymorphism observed for CsgA fibrils (39) supports both the β-helix model and the spine-based model. In the β-helix model, polymorphism may be averted as the entire protein is involved in structuring the fibril. In the spine-based model, polymorphism may be avoided due to the sequence specificity of nucleation. The gatekeeping residues in the R2-R4 repeats (44) might prevent segments within these repeats from serving as spines, thereby allowing for specific nucleation sites to mediate assembly of homogenous fibrils.

To obtain more insight on the structural properties of full-length CsgA fibrils we used ATR-FTIR spectroscopy. Previous analyses of amyloids (71, 85-88) revealed an FTIR signature spectral peak between 1611 and 1630 cm^−1^ while native β-sheet-rich proteins (71) show a peak at higher wavelengths of 1630–1643 cm^−1^. Often, a shift from higher to lower wavelengths is observed while monitoring fibril formation (86), which indicates the assembly of longer and planar sheets (71, 89), an increase in the number of β-strands in the β-sheet and/or the formation of stronger hydrogen bonding, typical of extremely stable amyloid fibrils (72, 86). In contrast to canonical amyloids, mature fibrils of the yeast prion HET-s(218-289) fragment, which was shown by solid-state NMR to form a β-solenoid-like fold (90), showed an FTIR peak at 1630-1631 cm^−1^ (91). This wavelength is at the threshold between wavelengths defining amyloids and native β-sheet proteins (71). CsgA fibrils show a peak at 1617 cm^−1^ (Fig. S7), which aligns with the range of wavelengths corresponding to rigid cross-β amyloid fibrils (71-73). An FTIR main peak at 1617 cm^−1^ was also shown for amyloid fibrils of apomyoglobin (92) and of γD-crystallin (93). Previous work showed a main FTIR peak of CsgA at 1623 cm^−1^, also within the lower wavelength range (47). Overall, the FTIR spectra of CsgA fibrils better support the formation of stable cross-β fibrils of in-register, tightly mated sheets. However, high-resolution structural work on full-length CsgA is needed to accurately describe the architecture of the fibrils.

### D-enantiomeric peptides designed against Alzheimer’s disease prevented biofilm development

Inspired by the structural resemblance of CsgA to human pathological amyloids, we found that two D-enantiomeric peptides, ANK6 and DB3DB3, originally designed to destabilize Aβ oligomers involved in Alzheimer’s disease (55-60), inhibit fibrillation of full-length CsgA, slowing its transition from an unstructured soluble configuration to insoluble fibrils (Figs. 2-3 and Figs. S9, S11&S12). These D-peptides also inhibited the fibrillation of spine segments from the R1 and R5 CsgA repeats (Fig. S10), supporting the role of these spines as critical elements in CsgA fibrillation. The ability of the D-peptides to mediate inhibition of both CsgA and Aβ provides further support for the structural similarity of their targets. Moreover, ANK6 and DB3DB3 did not affect the fibrillation of the cytotoxic *S. aureus* PSMα3, which forms cross-α amyloid-like fibrils (35), demonstrating specificity in structural properties targeted by the inhibitors.

The D-peptides ANK6 and DB3DB3 significantly reduce static biofilm formation of *Salmonella typhimurium* in a curli-dependent manner (Figs. 4-6 and Fig. S15) in agreement with their effects on CsgA fibrillation in-vitro. Similarly, other small molecules and peptidomimetics that have been shown to interfere with the in-vitro assembly of amyloids secreted by *E. coli*, *B. subtilis* and other bacteria prevent biofilm formation and pilus biogenesis (23, 77, 94-99). Here, we offer a novel strategy for disrupting curli amyloid formation, and prompt further investigation into the promiscuity of a class of anti-amyloid therapeutics as antivirulence agents targeting amyloid-structured biofilms.

### Can CsgA influence neurodegenerative diseases?

Amyloid patterns of protein folding and aggregation are highly conserved through evolution and appear in all kindgoms of life (100). Amyloids were even suggested to serve as prebiotic replication of information-coding molecules (101). The best known pathological manifestation of amyloid self-assembly is in association to neurodegenerative diseases, which involve the formation of transmissible self-propagating prion-like proteins (102, 103). Similarities in molecular structures between amyloids of different species may be involved in the creation of these prion-like agents through molecular mimicry (100), raising concerns regarding the exposure of humans to various food sources and microbes that contain amyloids (104-106). Indeed, microbial amyloids interact with the amyloids of host systems (23, 107), putatively providing some immune-evasive and survival strategies (31, 108), and were suggested to contribute to the pathology of aggregation diseases (107, 109-118). This might be related to the ability of seeds of amyloid fibrils from one species to nucleate monomers from another species (74, 119-121). Here we show that CsgA can cross-seed fibrillation of Alzheimer’s disease-associated Aβ (Fig. S8), in accordance with a previous report (74). A thought-provoking hypothesis is that curli-producing bacteria influence human neurodegenerative and aggregation diseases by nucleating, accelerating or deterring fibrillation of human amyloids. Another mechanism could be affecting the immune system, leading to inflammation and stress that are correlated with neurodegenerative diseases (122, 123). Future work is needed to characterize more microbial amyloids at the molecular and atomic level, study their inter-species interactions, and modulate their activities (38).

## Materials and Methods

### Peptides and reagents

Curli peptide segments (custom synthesis) at >98% purity were purchased from GL Biochem (Shanghai) Ltd. The segments used include LNIYQY, IYQYGG, VTQVGF, and TASNSS from CsgA (UniProt accession number P28307), and TAIVVQ from CsgB (UniProt accession number P0ABK7). The peptides were synthesized with unmodified termini for crystallography or with fully or semi capped termini (acetylated in the N-terminus and amidated in the C-terminus), as specified, for the other assays. The fibrillation inhibitors tested are peptides consisted of D-enantiomeric amino acids, which were C-terminally amidated. The sequences are as follows: ANK6: RKRIRLVTKKKR-NH2, DB3DB3: RPITRLRTHQNRRPITRLRTHQNR-NH2 and D3: RPRTRLHTHRNR-NH2. PSMα3 (custom synthesis) at >98% purity was purchased from GL Biochem (Shanghai) Ltd (UniProt accession number P0C805). Aβ_1-40_ and Aβ_1-42_ (custom synthesis) at >98% purity was purchased from GL Biochem (Shanghai) Ltd. D-peptides (custom synthesis) at >95% purity were purchased from either GL Biochem (Shanghai) Ltd, peptides&elephants (Potsdam, Germany), or JPT (Berlin, Germany). Thioflavin T (ThT), congo red and crystal violet were purchased from Sigma-Aldrich. Dimethyl-sulfoxide (DMSO) was purchased from Merck. Ultra-pure water was purchased from Biological Industries.

### Preparation of D-peptides

D-peptides inhibitors (ANK6, DB3DB3 and D3) were solubilized in ultra-pure water and their concentrations were calculated by spectrophotometer Nanodrop 2000c instrument (Thermo) using absorbance at 205 nm, with the specific extinction coefficient calculated for each peptide by the ‘protein parameter calculator’ (124) (http://nickanthis.com/tools/a205.html).

### Computational prediction of amyloid spine segments

Amyloidogenic propensities of segments from CsgA and CsgB were predicted using combined information from several computational methods, including ZipperDB (66), Tango (67, 68), Waltz (69) and Zyggregator (70).

### CsgA expression and purification

The protocol for CsgA expression and purification was adapted from Wang et al. (44). A plasmid containing CsgA sequence cloned into pET11d with a C-terminal His_6_-tag was kindly provided by the Chapman lab (University of Michigan, USA) (44). The plasmid was transformed to *E. coli* BL-21 cells grown overnight in 25 ml Luria-Bertani (LB) medium supplemented with 50 µg/ml ampicillin, and further diluted into 700 mL of the same medium incubated at 37°C with 220 rpm shaking until growth reached to OD_600_ of 0.8-0.9. CsgA expression was induced with 0.5 mM isopropyl β-D-1-thiogalactopyranoside (IPTG) for 1 hr. Bacterial cell pellets were harvested by centrifugation at 4,500 rpm for 25 min and stored at −20°C. Thawed cell pellets were resuspended in 25 ml of lysis buffer (8 M Guanidinium HCl, 50 mM potassium phosphate buffer pH 7.3) and incubated at room temperature (RT) with agitation for 18-24 hrs. The supernatant was separated using centrifugation at 10,000 g for 25 min and incubated with 1.6 ml of HisPur cobalt resin beads (Thermo scientific) equilibrated with lysis buffer at RT with agitation for 1 hr. The mixture was loaded on a disposable polypropylene column at 4°C and washed with 10 ml of 50 mM potassium phosphate buffer pH 7.3, followed by another column wash with the same buffer supplemented with 12.5 mM of Imidazole. The elution of CsgA was achieved with 125 mM imidazole in 50 mM potassium phosphate buffer pH 7.3. Freshly purified CsgA was filtered using a 30 kDa cut-off column at 4°C (Amicon® Ultra-4, Sigma-Aldrich) to remove insoluble protein aggregates and seeds. Imidazole was removed by desalting the protein solution at 4°C using Zeba spin 7K desalting column (ThermoFisher Scientific) into 50 mM potassium phosphate buffer pH 7.3. CsgA concentration was determined by measuring absorption at 280 nm calculated by Molar extinction coefficient of 11,460 M^−1^ cm^−1^, as determined via the Expasy server (http://web.expasy.org/cgi-bin/protparam/protparam). The identity of CsgA was confirmed by Western Blot using Anti-6X His tag antibody (not shown).

### Multi angle light scattering size exclusion chromatography analysis to determine oligomerization state

Multi angle light scattering size exclusion chromatography (SEC-MALS) analysis of freshly purified CsgA was performed to determine the accurate molecular weight and the oligomerization state of soluble CsgA. CsgA was concentrated to 1.8 mg/ml using a 3 kDa cut-off spin column at 4ºC (Amicon® Ultra-4, Sigma-Aldrich). The size exclusion chromatography (SEC) was performed over a size-exclusion column (Superdex 75 10/300) operated by AKTA avant. The multi angle light scattering (MALS) was performed with miniDAWN TREOS (WYATT Technology) and its companion Optilab T-rEX (WYATT Technology) dRI detector. The characterization and analysis of the SEC-MALS results was done using ASTRA software (WYATT Technology).

### Fibrillation assays

#### Transmission Electron Microscopy (TEM) visualizing CsgA fibrils

TEM was performed to visualize CsgA fibrils, and to test the effect of the D-peptide inhibitors. CsgA was purified as described above and incubated at a concentration of 30 µM overnight at 25°C with 300 rpm shaking in a plate reader, with and without 300 µM D-peptides (1:10 ratio) before fixation. Five-microliter samples were applied directly onto copper TEM grids with support films of Formvar/Carbon (Ted Pella), which were charged by glow-discharge (PELCO easiGlow, Ted Pella) immediately before use. Grids were allowed to adhere for 2 min and negatively stained with 5 μl of 2% uranyl acetate solution. Micrographs were recorded using FEI Tecnai G2 T20 S-Twin transmission electron microscope at an accelerating voltage of 200 KeV located at the Department of Materials Science & Engineering at the Technion, Israel, or using FEI Tecnai T12 G2 transmission electron microscope operated at an accelerating voltage of 120 kV located at the Ilse Katz Institute for Nano-Science and Technology at Ben-Gurion University of the Negev, Israel.

#### Thioflavin T kinetic assays of CsgA

Thioflavin T (ThT) is a widely used “gold standard” stain for identifying and exploring formation kinetics of amyloid fibrils, both in-vivo and in-vitro. Fibrillation curves in presence of ThT commonly show a lag time for the nucleation step, followed by rapid aggregation. Fibrillation kinetics of CsgA with and without D-peptide inhibitors was monitored using this approach. Freshly purified CsgA in 50 mM potassium phosphate buffer pH 7.3 was mixed with filtered ThT from stock made in ultra-pure water diluted in the same buffer, and with D-peptides from stock made in ultra-pure water. Final concentrations in the reaction wells were 15 µM CsgA mixed with 0, 15, 75 or 150 µM D-peptides, and 20 µM ThT in a final volume of 100 µl. The control wells included all the components except the D-peptides that were replaced with pure water at the same volume. The reaction mixture was carried out in a black 96-well flat-bottom plate (Greiner bio-one) covered with a thermal seal film (EXCEL scientific) and incubated in a plate reader (CLARIOstar, BMG Labtech) at a temperature of 25°C with orbital shaking at 300 rpm for 30 sec before each measurement. ThT fluorescence was recorded every two minutes using an excitation of 438±20 nm and an emission of 490±20 nm. The measurements were conducted in triplicates, and the entire experiment was repeated at least three times.

#### Pretreatment of Aβ peptides and preparation of CsgA fibril seeds for ThT kinetics

Fibrillation of Aβ_1-42_ and Aβ_1-40_ was monitored using ThT kinetics with and without the addition of CsgA fibril seeds. Preparation of CsgA fibril seeds was performed as previously described (119). Specifically, CsgA fibrils, obtained after incubation of freshly purified CsgA for one day at the indicated concentrations (0.2-2 μM), were sonicated with a VC750 VibraCell sonicator (sonics, CT, USA) using 30% amplitude with three 15 sec bursts on ice, 50 sec apart.

The Aβ peptides were pretreated in order to dissolve potential aggregates in the lyophilized powder. Aβ_1-40_ was dissolved to 1 mg/ml in 1,1,1,3,3,3- hexafluoro-2-propanol (HFIP), bath sonicated at RT for 10 min, incubated for 1 hr at 37°C, and dried in the chemical hood for three days. Residual HFIP was dried using SpeedVac (RVC 2-18 CDplus, CHRIST). Aβ_1-42_ was pretreated as previously described (119). Specifically, Aβ_1-42_ was dissolved to 1 mg/ml in 100% Trifluoroacetic acid (TFA), bath sonicated at RT for 10 min and dried using SpeedVac for 1 hr. The pellet was dissolved in 1 ml HFIP and dried using SpeedVac for 1 h. The HFIP process was repeated three times. Aβ_1-42_ and Aβ_1-40_ were stored in *-*20°C until use. The ThT kinetic assay was carried out as described above with 50 μM Aβ, 20 μM ThT, and with or without the CsgA fibril seeds. The measurements were performed in triplicates, and the entire experiment was repeated at least three times with similar results.

#### Thioflavin T kinetic and TEM assay of CsgA spine segments

Capped CsgA segments with capped termini (acetylated in the N-terminus and amidated in the C-terminus) were used to mimic their chemical nature in the full-length protein. For TEM visualization, the peptides were dissolved to 1 mM in DMSO and incubated at 37ºC with 300 rpm shaking for few days. TEM grids were prepared and visualized as described above. For the ThT assay, the peptides were dissolved in DMSO to 10 mM stocks and kept at −20ºC until usage. The peptides were thawed and diluted directly into the reaction buffer made of 50 mM potassium phosphate buffer pH 7.3 mixed with filtered ThT from stock made in ultra-pure water. The final concentrations were 100, 150, 200 or 500 µM of CsgA segments and 20 µM ThT in a final volume of 100 µL per well. The ThT reaction was conducted as described above. The measurements were performed in triplicates, and the entire experiment was repeated at least twice with similar results.

#### Thioflavin T kinetic assay of CsgA spine segments in the presence of the D-peptides

CsgA segments capped at the C-terminus were dissolved in DMSO to 10 mM stocks and kept at −20ºC until usage. The peptides were thawed and diluted directly into the reaction buffer made of 50 mM potassium phosphate buffer pH 7.3 mixed with filtered ThT from stock made in ultra-pure water, and with D-peptides from stock made in ultra-pure water to the indicated final concentrations. The final concentration of ThT was 20 µM ThT in a final volume of 100 µL per well. The reaction was conducted as described above. The measurements were performed in triplicates, and the entire experiment was repeated at least three times with similar results.

#### Thioflavin T kinetic assay of PSMα3 with D-peptides

Fresh ThT reaction solution was prepared by a 10-fold dilution of a filtered ThT stock, made in ultra-pure water, into a reaction buffer containing 10 mM sodium phosphate buffer and 150 mM NaCl at pH 8.0. Lyophilized powder of PSMα3 was freshly dissolved in TFA-HFIP (1:1) to a concentration of 5 mg/ml, sonicated for 3-5 min in a sonication bath, and evaporated using SpeedVac. The pretreated peptide was dissolved to 1 mM, on ice, in ultra-pure water, and sonicated for 3-5 min using a sonication bath. The peptide was then diluted, on ice, with a filtered reaction buffer, and centrifuged for 5 min at 4°C and 10,000 rpm. The D-peptides DB3DB3 and ANK6 were each dissolved in ultra-pure water to 50 mM stock concentration and stored at −80°C until used. The reaction was carried out in a Greiner bio-one black 96 well flat-bottom plate. For each reaction well, the supernatant of the PSMα3 solution was separately supplemented with the D-peptides in 1:0 or 1:5 PSMα3:D-peptide molar ratios, and supplemented with ThT to final concentrations of 50 µM PSMα3, 200 µM ThT and 0 or 250 µM D-peptide. For blank subtractions, ThT was mixed with the reaction buffer alone, or with 250 µM of each of the D-peptides separately. The reaction plate was immediately covered with a silicone sealing film (ThermalSeal RTS), and incubated in a plate reader (CLARIOstar) at 37ºC, with 500 rpm shaking for 15 sec before each cycle, and up to 1000 cycles of 5 min each. Measurements were made in triplicates. ThT fluorescence at an excitation of 438±20 nm and emission of 490±20 nm was recorded. All triplicate values were averaged, appropriate blanks were subtracted, and the resulting values were plotted against time. Calculated standard errors of the mean are presented as error bars.

### CsgA fibrillation inhibitors tested using SDS-PAGE migration

CsgA fibrils are insoluble in sodium dodecyl sulfate (SDS), even after boiling, thus are unable to migrate into SDS polyacrylamide gel electrophoresis (SDS-PAGE), in contrast to soluble CsgA (77). Therefore, this method can be used to monitor CsgA fibrillation and test potential inhibitors. Freshly purified recombinant CsgA was diluted in 50 mM potassium phosphate buffer, pH 7.3, mixed with diluted D-peptides into 1 mL tubes. Final concentrations used were 10 μM CsgA and 50 μM D-peptides (1:5 molar ratio). Samples were then incubated for 24 hrs at 25ºC with 300 rpm shaking. 20 μL from the samples were then mixed with 10 μL 3× SDS sample buffer with dithiothreitol (DTT) and incubated at 95°C for 10 min. Samples were loaded on a 15% SDS-PAGE gel for protein migration, and a known concentration of BSA solution was added to quantify the amount of protein in each band. Coomassie Stain (Expedeon - InstantBlue Coomassie Protein Stain) was performed to visualize the migration of soluble CsgA in the gel. Imaging was performed using gel documentation system (Gel Doc – BioRad).

### Circular dichroism

CsgA fibrillation involves conformational changes from an unstructured coil into a β-rich conformation. We used circular dichroism (CD) in order to monitor the secondary structure transitions of CsgA alone and in the presence of the D-peptide inhibitor ANK6. Immediately prior to the CD experiment, freshly purified recombinant CsgA (in 50 mM potassium phosphate buffer, pH 7.3) was dialyzed at 4°C using a Zeba spin 7K desalting column (ThermoFisher Scientific) to 2 mM potassium phosphate buffer, pH 7.3, to reduce background signal during CD measurements. CsgA was directly diluted into the CD cuvette to a final concentration of 9 µM. At the same time, 9 µM CsgA was mixed in a second cuvette with 45 µM ANK6 (1:5 molar ratio) diluted from 50 mM stock made in ultra-pure water. The signal from blank solutions of either 2 mM potassium phosphate buffer pH 7.3 alone, or of the same buffer, containing 45 µM of ANK6, were taken at the beginning of the experiment, just before the addition of CsgA into the appropriate cuvette. CD measurements were performed at several time point over 18 hours, with cuvettes being incubated at RT between measurements, and mixed thoroughly before each measurement.

Far UV CD spectra were recorded on the Applied Photophysics PiStar CD spectrometer (Surrey, UK) equilibrated with nitrogen gas, using a 0.1 mm path-length demountable quartz cell (Starna Scientific, UK). Changes in ellipticity were followed from 290 nm to 180 nm with a 1 nm step size and a bandwidth of 1 nm. The measurements shown are an average of three scans for each time point, or two scans for blanks, captured at a scan rate of 1 second per point.

### Attenuated total internal reflections Fourier transform infrared spectroscopy

Freshly purified recombinant CsgA in 50 mM potassium phosphate buffer pH 7.3 was frozen in liquid nitrogen and lyophilized over-night to complete dryness. The dry CsgA powder was dissolved in deuterium oxide (D_2_O) to remove water residues interrupting the FTIR signal, and further lyophilized. This procedure was repeated twice. The treated sample was then dissolved in D_2_O to 20 mg/ml immediately prior to measurements. Five μl sample of CsgA solution was spread on the surface of the ATR module (MIRacle Diamond w/ZnSe lens 3-Reflection HATR Plate; Pike Technologies) and let dry under nitrogen gas to purge water vapors. Absorption spectra were recorded on the dry fibrils using a Tensor 27 FTIR spectrometer (Bruker Optics). Measurements were performed in the wavelength range of 1500-1800 cm^−1^ in 2 cm^−1^ steps and averaged over 100 scans. Background (ATR crystal) and blank (D_2_O) were measured and subtracted from the final spectra. The amide I’ region of the spectra (1600 – 1700 cm^−1^) is presented in the graphs.

### Crystallization of CsgA segments

Peptides synthesized with free (unmodified) termini were used for crystallization experiments to facilitate crystal contacts. IYQYGG at 10 mM was dissolved in water. VTQVGF at 10 mM was dissolved in 100% DMSO. LNIYQY crystals grow in a mixture of 10 mM LNIYQY and 10 mM TAIVVQ (a segment from CsgB) dissolved in 82% DMSO. TASNSS crystals grow in a mixture of 30 mM TASNSS and 10 mM TAIVVQ dissolved in 82% DMSO. Peptide solution drops at 100 nL volume were dispensed by the Mosquito automated liquid dispensing robot (TTP Labtech, UK) located at the Technion Center for Structural Biology (TCSB) to crystallization screening plates. Crystallization experiments were set up using the hanging drop method in 96-well plates with 100 µL solution in each well. The drop volumes were 150-300 nL. All plates were incubated in a Rock imager 1000 robot (Formulatrix) at 293 °K. Micro-crystals grew after few days and were mounted on glass needles glued to brass pins(125). No cryo protection was used. Crystals were kept at RT prior to data collection. Reported structures were obtained from drops that were a mixture of peptide and reservoir solution as follows: IYQYGG: 10 mM IYQYGG, 0.1 M sodium acetate pH 4.6, and 2.0 M sodium formate. LNIYQY: 10 mM LNIYQY, 10 mM TAIVVQ, 0.1 M HEPES pH 7.5, and 20%v/v Jeffamine M-600. VTQVGF: 10 mM VTQVGF, 3.0 M sodium chloride, 0.1 M BIS-Tris pH 5.5. TASNSS: 30 mM TASNSS, 10 mM TAIVVQ, 0.2 M lithium sulfate, 0.1 M Tris-HCl pH 8.5, and 30% (w/v) polyethylene glycol 4000.

### Structure determination and refinement

X-ray diffraction data was collected at 100°K using 5° oscillation. The X-ray diffraction data for VTQVGF was collected at the micro-focus beamline ID23-EH2 of the European Synchrotron Radiation Facility (ESRF) in Grenoble, France; wavelength of data collection was 0.8729 Å. The X-ray diffraction data for LNIYQY, IYQYGG and TASNSS was collected at the micro-focus beamline P14 operated by EMBL at the PETRAIII storage ring (DESY, Hamburg, Germany); wavelength of data collection was 0.9763 Å. Data indexation, integration and scaling were performed using XDS/XSCALE (126). Molecular replacement solutions for all segments were obtained using the program Phaser (127) within the CCP4 suite (127-129). The search models consisted of available structures of geometrically idealized β-strands. Crystallographic refinements were performed with the program Refmac5 (129). Model building was performed with Coot (130) and illustrated with Chimera (131). There were no residues that fell in the disallowed region of the Ramachandran plot. Crystallographic statistics are listed in Table S1.

### Calculations of structural properties

The Lawrence and Colman’s shape complementarity index (132) was used to calculate the shape complementarity between pairs of sheets forming the dry interface (Table S2). The surface area buried was calculated with Chimera (UCSF) with default probe radius and vertex density are 1.4 Å and 2.0/Å^2^, respectively. The number of solvent accessible surface area buried is the average area buried of one strand within two β-sheets (total area buried from both sides is double the reported number in Table S2)

### Bacterial strains and growth conditions

*Salmonella typhimurium* bacterial strains used: the MAE52 strain display a constitutive multicellular morphotype mediated by the expression of the *agfD* operon leading to the production of an adhesive extracellular matrix consisting of curli fimbriae (previously called thin aggregative fimbriae (*agf*)) and cellulose (133). The MAE150 (∆*bcsA*) strain sustained a deletion of the bacterial cellulose synthesis (*bcs*), a gene encoding cellulose synthase (134). *S*. *typhimurium* bacterial MAE52 and MAE150 strains were grown on LB agar plates without NaCl at 30°C. For liquid media growth a single bacterial colony was picked from an agar plate and dipped in LB media (lacking NaCl) followed by incubation at 30°C with vigorous shaking.

### Static biofilm analysis

The evaluation of static biofilm production was adapted from a previous study (135). Bacterial suspension of *S*. *typhimurium* MAE52 with OD_600_ of 0.05-0.13 were incubated in 200 µl LB media containing D-peptide inhibitors at increasing concentrations (0, 5, 10, 20, or 40 µM) for 48 hours at 30°C in a 96-well plate (nunclon, Roskilde, Denmark). The plate was then washed twice with 0.9% saline (w/v).

#### Surface attached biofilm assay

The protocol was adapted from previous reports (136, 137). Specifically, after washing the plates, biofilm cells that remained attached to the bottom of the micro-wells were fixated with 4% paraformaldehyde (w/v), washed tree times with 0.9% saline (w/v), stained with propidium iodide and analyzed using Zeiss confocal laser scanning microscopy (CLSM) with 10 × lenses and 522 nm and 682 nm excitation/emission filters. All signals were recorded using Zeiss 710 microscopy (Jena, Germany) located at the Technion, Israel.

#### Total biofilm biomass assay

In a separated experiment, pellicle biofilm attached to the sides of the micro-wells at the liquid-air interface as well as the surface-attached biofilm were analyzed via standard crystal violet staining. After washing the wells, as described, 200 µl of 0.1% of crystal violet staining solution was added to each well and incubated for 15 minutes followed by removal of the dye and washing with 0.9% saline. A 200 µl of 95% ethanol solution was added to each micro-well and incubated for 30 minutes after which 100 µl from each well were transferred to a new 96-micro-well plate and recorded at absorbance of OD_600_ with a Synthergi HT (Biotek, Winooski, USA) device.

### Curli production analysis using congo red staining

Curli (along with cellulose) production is associated with CR staining in *S*. *typhimurium* (79), thus the lack of CR adsorption by the bacterial cells is evident in the residual agar below the colony (25). *S*. *typhimurium* MAE150 was grown overnight in LB media and then diluted to obtain a bacterial suspension with OD_600_ of 0.05-0.13. D-peptide inhibitors at increasing final concentrations (0, 10, 20, or 40 µM) were added to the media and 3 µl of the suspension was spotted on a LB-agar plate supplemented with 20 µg/ml CR. The bacterial biofilm colonies were allowed to grow for 48 h at 30 °C, after which the colony was immersed in 0.4% paraformaldehyde followed by the removal of the colony with double distilled water washes. In order to test the amount of CR that remained on the agar after the removal of the colonies, the agar was dissolved as previously described (135). In brief, the agar underneath the removed colony was cut out, and incubated at 55 °C for 10 min with 1 ml of binding buffer solution (hylabs, Rehovot, Israel). The CR residual staining in the melted agar was measured by absorbance at OD_510_ (optical dentistry was calibrated after scanning the optimum wave length for CR detection) using nanodrop 2000c (Thermo).

### Statistical analysis

Statistical analyses were carried out for the total biofilm biomass assay with crystal violet staining and for curli production analysis using CR staining. The data were not normally distributed according to the Shapiro Wilk test. The analyses were performed using the non-parametric Kruskal-Wallis test for comparing three independent groups, the Mann-Whitney non-parametric test for comparing two independent groups (all variables calculated have small sample size (N ≥10)). All tests applied were two-tailed, and a p-value of 5% or less was considered statistically significant. Analyses were performed using SPSS software, version 24.

## Data availability

The raw data that support the findings of this study are available on request from the corresponding author. Coordinates and structure factors for the X-ray crystal structures have been deposited in the protein data bank (PDB) with accession codes 6G8C (IYQYGG), 6G8D (LNIYQY), 6G8E (VTQVGF) and 6G9G (TASNSS).

## Acknowledgments

We are grateful to Prof. Sima Yaron for supplying the bacterial strains used in this study as well as the research space for conducting the experiments. We thank O. Tabachnikov, M. Chapman, and N. Jain, for help with experiments. We acknowledge Y. Pazy-Benhar and D. Hiya at the Technion Center for Structural Biology (TCSB), and the Life Sciences and Engineering Infrastructure Center at the Technion. We acknowledge support from Yaron Kauffmann from the MIKA electron microscopy center of the department of material science & engineering at the Technion, the Russell Berrie Electron Microscopy Center of Soft Matter at the Technion, Israel, and the service provided by Einat Nativ-Roth and Alexander Upcher from the Ilse Katz Institute for Nano-Science and Technology at Ben-Gurion University of the Negev, Israel. This research was supported by the I-CORE Program of the Planning and Budgeting Committee and The Israel Science Foundation, Center of Excellence in Integrated Structural Cell Biology (grant no. 1775/12), DFG: Deutsch-Israelische Projektkooperation (DIP) (grant no. LA 3655/1-1), Israel Science Foundation (grant no. 560/16), University of Michigan – Israel Collaborative Research Grant, and BioStruct-X, funded by FP7. The synchrotron MX data collection experiments were performed at beamline ID23-EH2 at the European Synchrotron Radiation Facility (ESRF), Grenoble, France and at beamline P14, operated by EMBL Hamburg at the PETRA III storage ring (DESY, Hamburg, Germany). We are grateful to the teams at ESRF and EMBL Hamburg. We would like to thank Thomas R. Schneider and Gleb Bourenkov for the assistance in using the P14 beamline.

## Supporting Information

**Figure S1.**
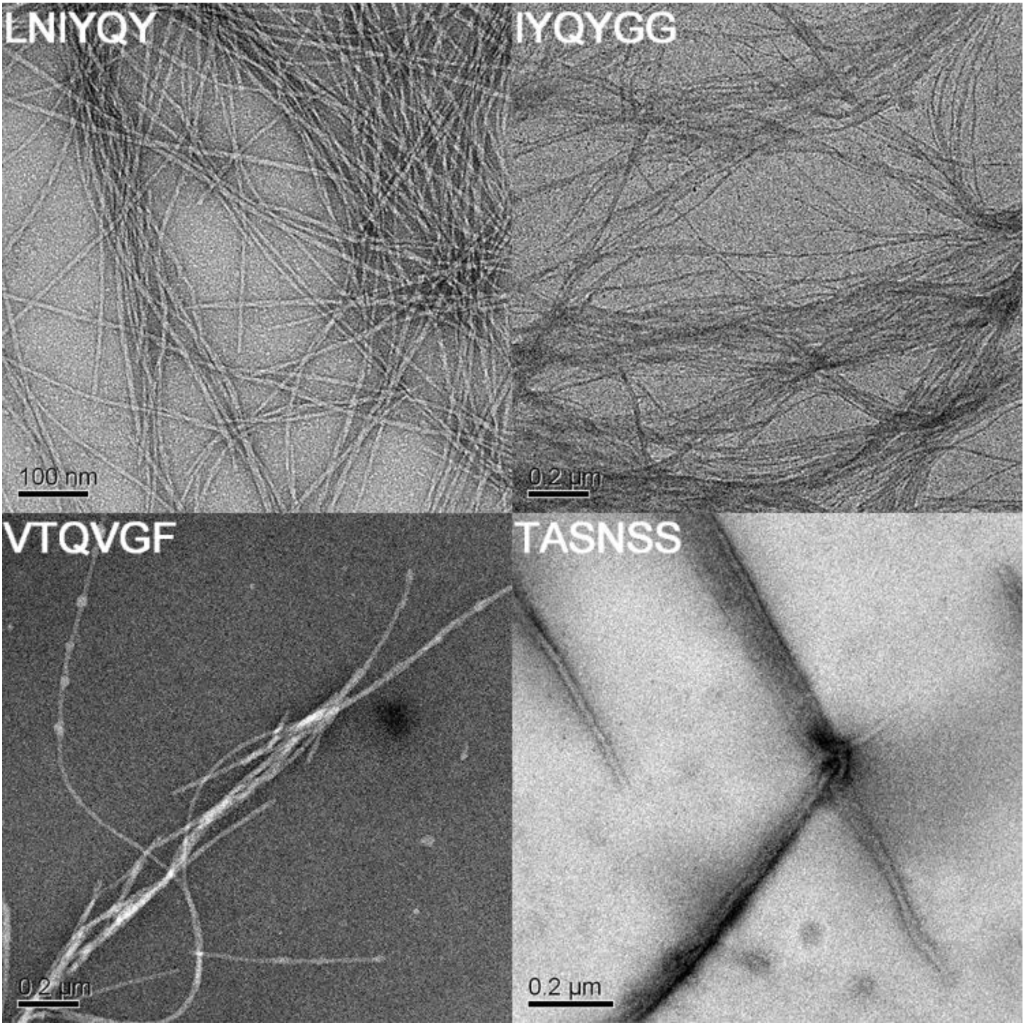
Electron micrographs of fibrils of CsgA segments. TEM micrographs visualizing fibrils of _45_LNIYQY_50_ (R1), _47_IYQYGG_52_ (R1), _137_VTQVGF_142_ (R5) and _129_TASNSS_134_ (R4-R5 loop). Scale bars are indicated (100 nm for _45_LNIYQY_50_ and 200 nm for the others).

**Figure S2.**
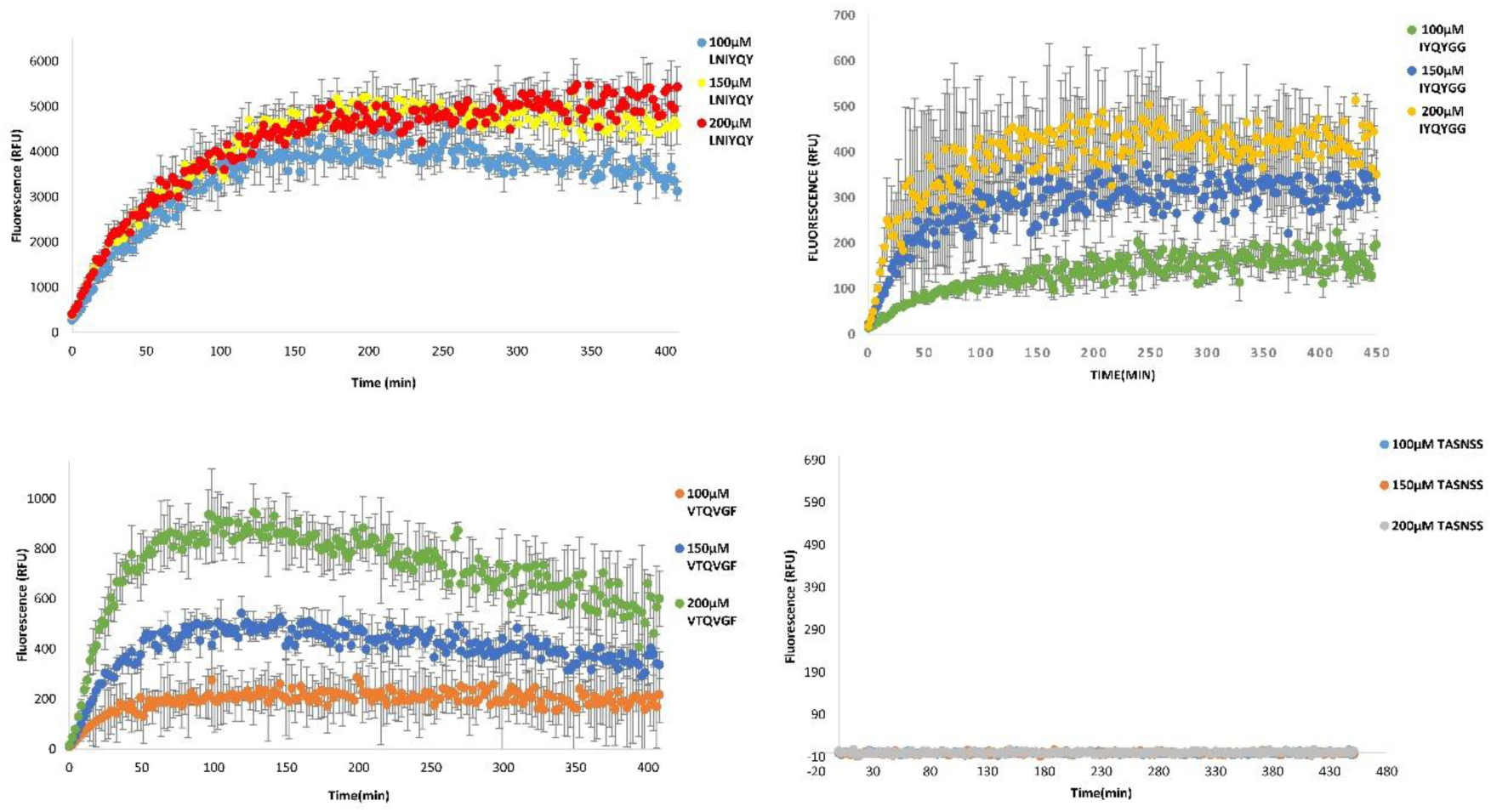
Concentration-dependent ThT fibrillation kinetics of CsgA segments. The graphs represent averaged fluorescence reading of ThT triplicated measurements of the CsgA segments at 100, 150 and 200 µM. Error bars represent standard error of the mean calculated from a triplicate.

**Figure S3.**
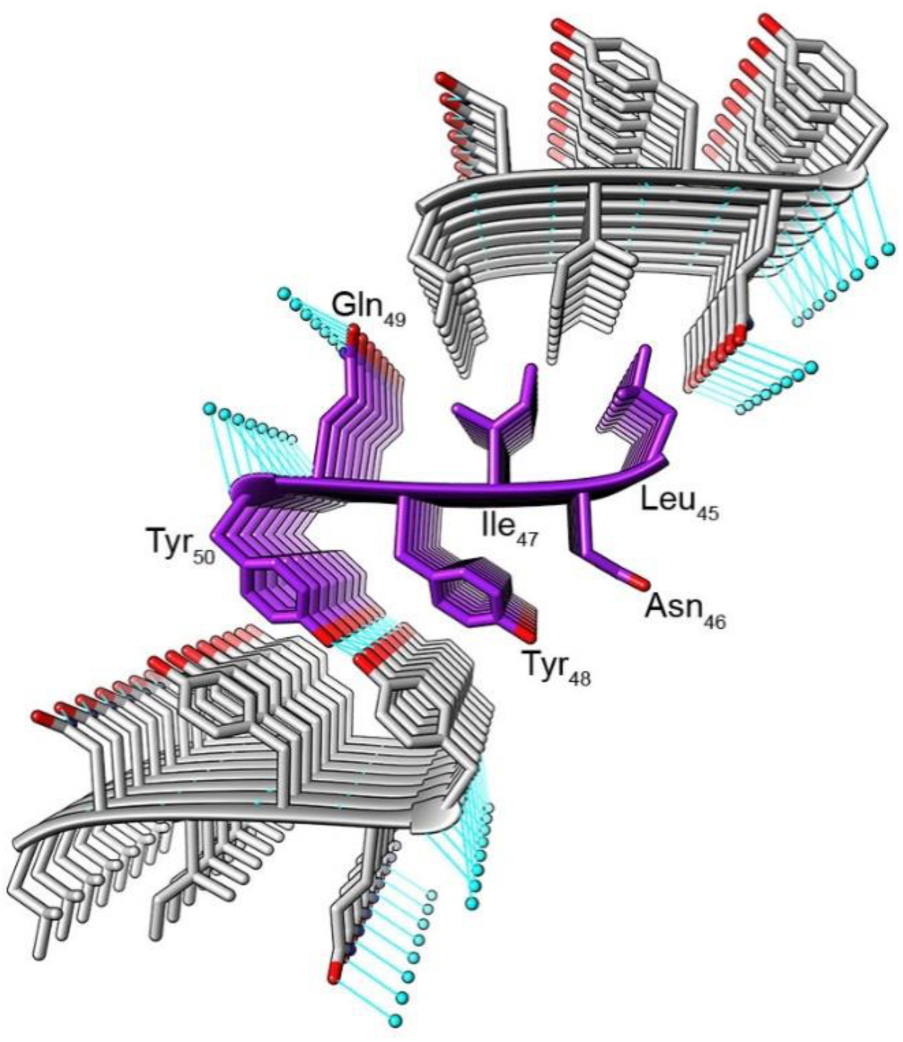
Structural description of the _45_LNIYQY_50_ fibril. The crystal structure of _45_LNIYQY_50_ demonstrates the formation of a cross-β steric zipper fibril composed of mated, parallel β-sheets. Two possible tight and dry interfaces are observed in the crystal. The first dry interface between mated β-sheets is mostly hydrophobic, formed between facing and tightly packed Leu45 and Ile47 residues flanked by Gln49 side chains. In this conformation, water molecules running along the fibril axis may form hydrogen bonds with the Gln49 side chains as well as with the C-terminus carboxyl group. The second interface is predominantly mediated by two tyrosine residues (Tyr48 and Tyr50). These tyrosine residues face each other, forming a tight and dry interface along the fibril axis. Tyr50 from each strand may form hydrogen bonds with equivalent tyrosines from facing and adjacent strands, creating a network of hydrogen bonds within the dry interface along the fibril axis. The Asn46 residues are facing the same direction as the tyrosines on the β-strands, but do not directly participate in the interface between mating sheets. However, these asparagine residues putatively form a ladder of hydrogen bonds along the fibril axis (not shown), further stabilizing the fibril structure. The carbons of each β-sheet are colored either gray or purple; heteroatoms are colored by atom type (nitrogen in blue, oxygen in red). Water molecules are shown as small cyan spheres. Hydrogen bonds are shown in cyan lines.

**Figure S4.**
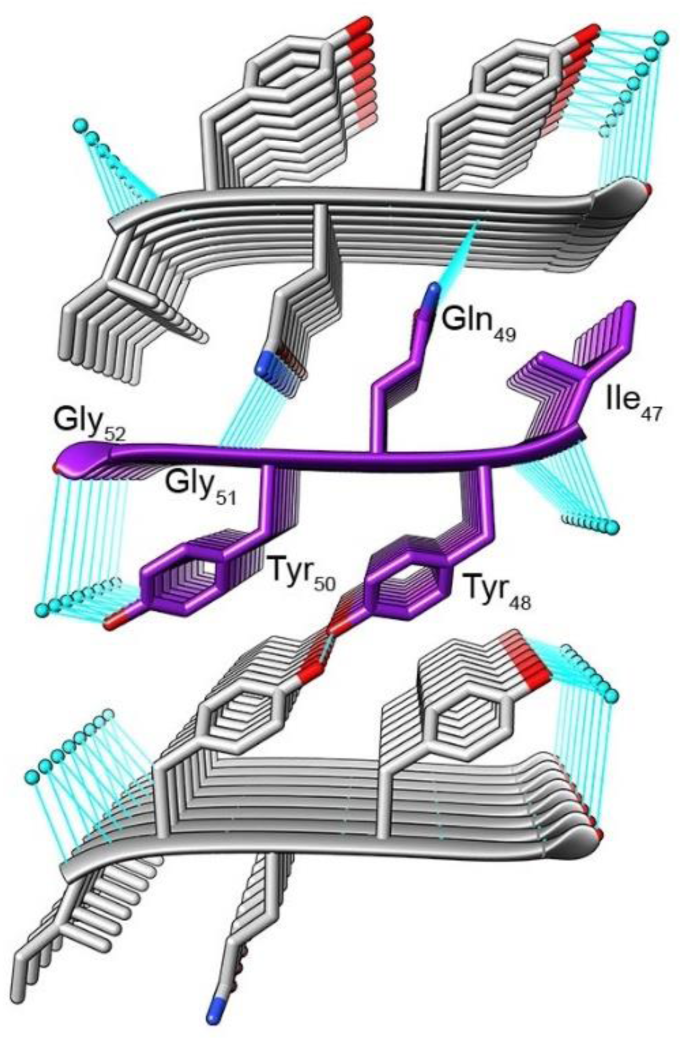
Structural description of the _47_IYQYGG_52_ fibril. The _47_IYQYGG_52_ segment, which partially overlaps with _45_LNIYQY_50_, also forms two possible dry zipper interfaces. The first interface is mediated via Ile47, Gln49, and Gly51 from both sides of the mated β-sheets. Each Gln49, located in the middle of the interface, may participate in hydrogen bonds with adjacent glutamines along the sheet (not shown) and with the backbone oxygen of Tyr50. As with _45_LNIYQY_50_, the second interface is mediated by Tyr48 and Tyr50. However, in _47_IYQYGG_52_, Tyr48 from each strand forms hydrogen bonds with equivalent tyrosines from facing and adjacent strands, creating a network of hydrogen bonds within the dry interface along the fibril axis. Water molecules flank the dry interface, putatively engaging in hydrogen bonds with Tyr50, with the C-terminus carboxyl group, and with the N-terminal amine group along the fibril axis. Coloring scheme is as in Fig. S3.

**Figure S5.**
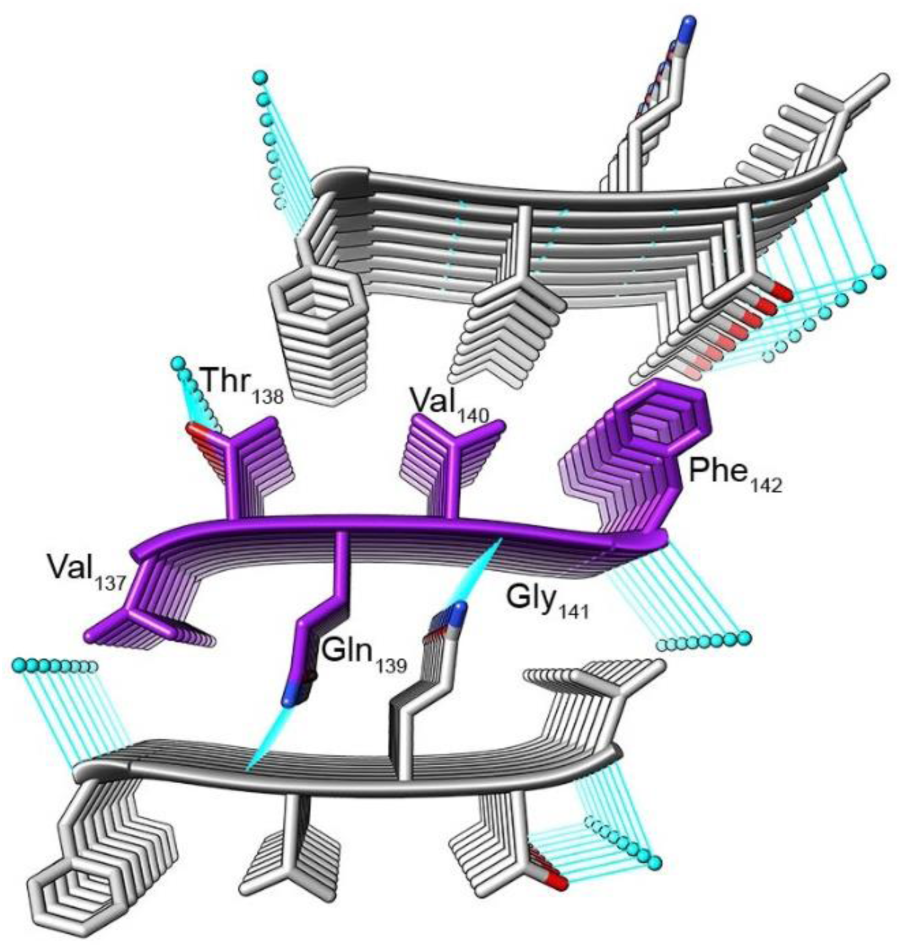
Structural description of the _137_VTQVGF_142_ fibril. The crystal structure of _137_VTQVGF_142_ shows two possible dry interfaces between parallel mated β-sheets. One interface is mediated by Thr138, Val140, and Phe142. These residues are tightly packed forming a hydrophobic, dry, interface, with the side chain oxygen of Thr138 positioned at the periphery of the interface, forming putative hydrogen bonds with water molecules along the fibril axis. The second dry interface is mediated via Val137, Gln139, and Gly141. As with _47_IYQYGG_52_, the glutamines are located in the middle of the interface and engage in putative hydrogen bonds with adjacent glutamines along the sheet (not shown) as well as with backbone oxygens, here of Val140. Coloring scheme is as in Fig. S3.

**Figure S6.**
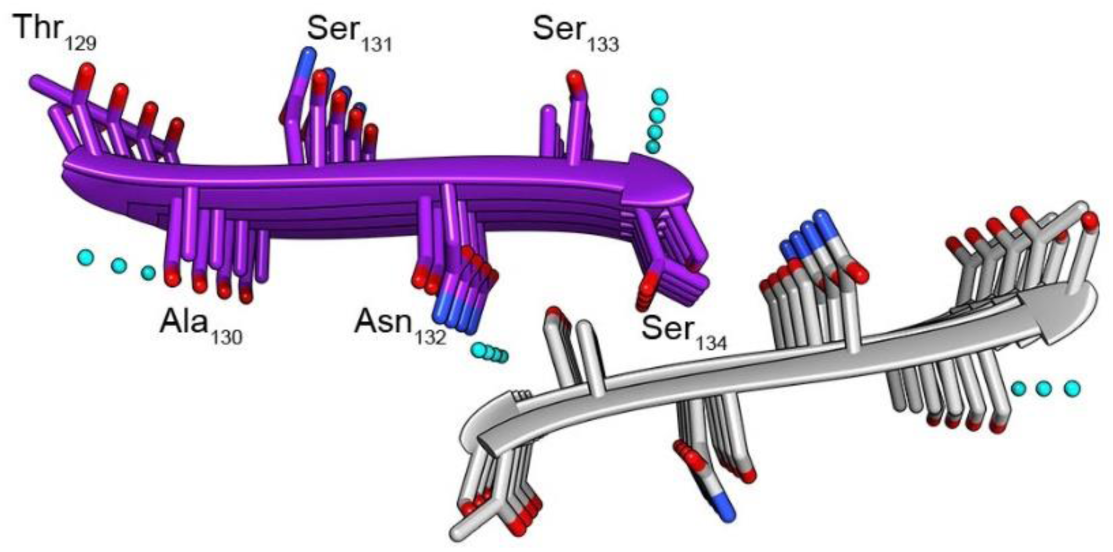
Structural description of the _129_TASNSS_134_ fibril. _129_TASNSS_134_ from the R4-R5 loop region was selected as a control sequence. This segment was predicted by computational methods to be amyloidogenic but is located in a region not implicated in fibrillation. In contrast to the other three segments that form tightly packed steric zipper structures, the _129_TASNSS_134_ segment forms extended chains yielding anti-parallel β-sheets. Each β-sheet is composed of anti-parallel strands putatively stabilized within the sheet both by hydrogen bonds between backbone atoms along the sheets as well as electrostatic interactions between the C-and N-termini. Furthermore, the C-terminal Ser134 can form hydrogen bonds with the N-termini of adjacent strands on the same sheet. In contrast to the other three spine segments from the R1 and R5 repeats, the β-sheets of _129_TASNSS_134_ do not mate via a tight interface. Each sheet is not directly facing another sheet but shifted. Nevertheless, several inter-sheet interactions stabilize this configuration, including possible hydrogen bonds between Thr129 and Ser133, Ser134 and the backbone oxygen of Asn132, and Ser131 and the N-terminus (bonds not shown due to antiparallel orientation that prevents a clear visualization). This architecture is chemically stable though it does not strictly belong to a class of steric zippers. In accordance with its unusual structure, this segment forms ribbon-like structures with atypical morphology as demonstrated by TEM (Fig. S1). These atypical ribbons do not bind ThT (Fig. S2). Coloring scheme is as in Fig. S3.

**Figure S7.**
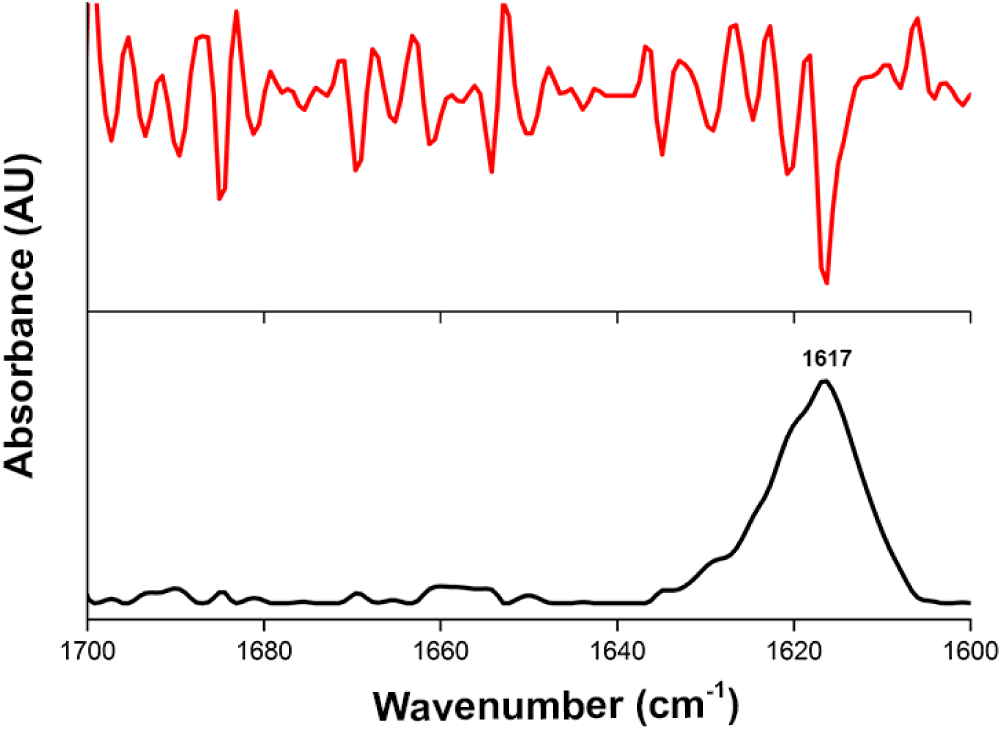
ATR-FTIR spectra demonstrates the cross-β architecture of full-length CsgA fibrils. Attenuated total internal reflection Fourier transform infrared (ATR-FTIR) spectroscopy of the amide I’ region (1600-1700 cm^−1^) of CsgA fibrils shows a main peak at 1617 cm^−1^ corresponding to rigid amyloid fibrils (1-3). The black line represents the ATR spectra and the red line is the second derivative.

**Figure S8.**
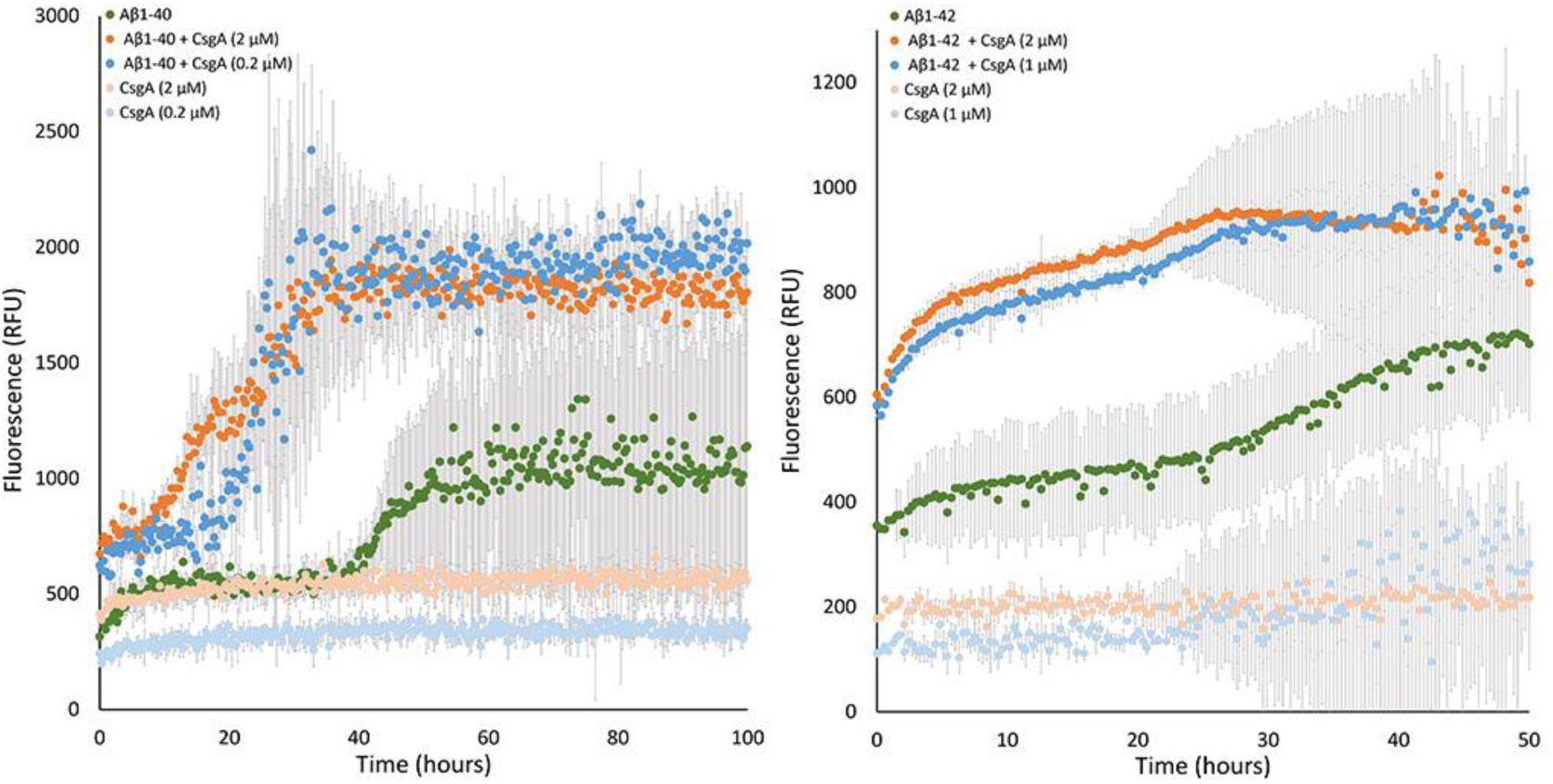
Seeds of CsgA fibrils enhance fibrillation of Aβ_1-40_ and Aβ_1-42_. The graphs represent averages of triplicate ThT measurements of 50 µM Aβ_1-40_ (**A**) or Aβ_1-42_ (**B**) alone and with the addition of seeds of CsgA fibrils. The seeds were prepared by sonication of CsgA fibrils formed after a 24 hrs incubation of freshly purified recombinant CsgA at different initial concentrations (0.2 or 2 µM (**A**), and 1 or 2 µM (**B**)). CsgA fibril seeds alone were used as a control and showed minor ThT binding in accordance with their low concentration. Error bars represent standard error of the mean calculated from triplicate measurements. Each graph represents three independent experiments repeated on different days.

**Figure S9.**
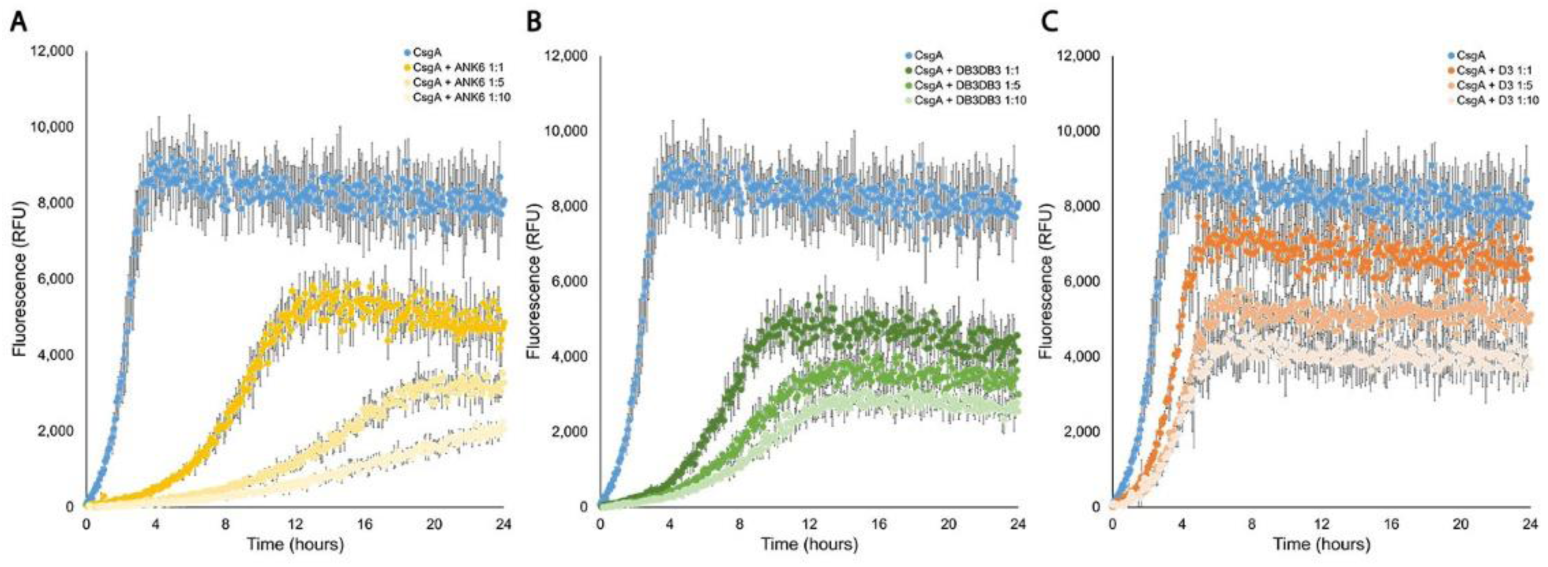
D-peptides inhibit fibrillation of CsgA in dose-dependent manners. The graphs show mean fluorescence readings of triplicate ThT measurements of CsgA with or without ANK6 (**A**), DB3DB3 (**B**) or D3 (**C**) at different molar ratios as shown in the color-coded bar. Error bars represent standard error of the mean calculated from triplicates. The D-peptides delayed fibril formation of CsgA and reduced the fluorescence signal in dose-dependent manners.

**Figure S10.**
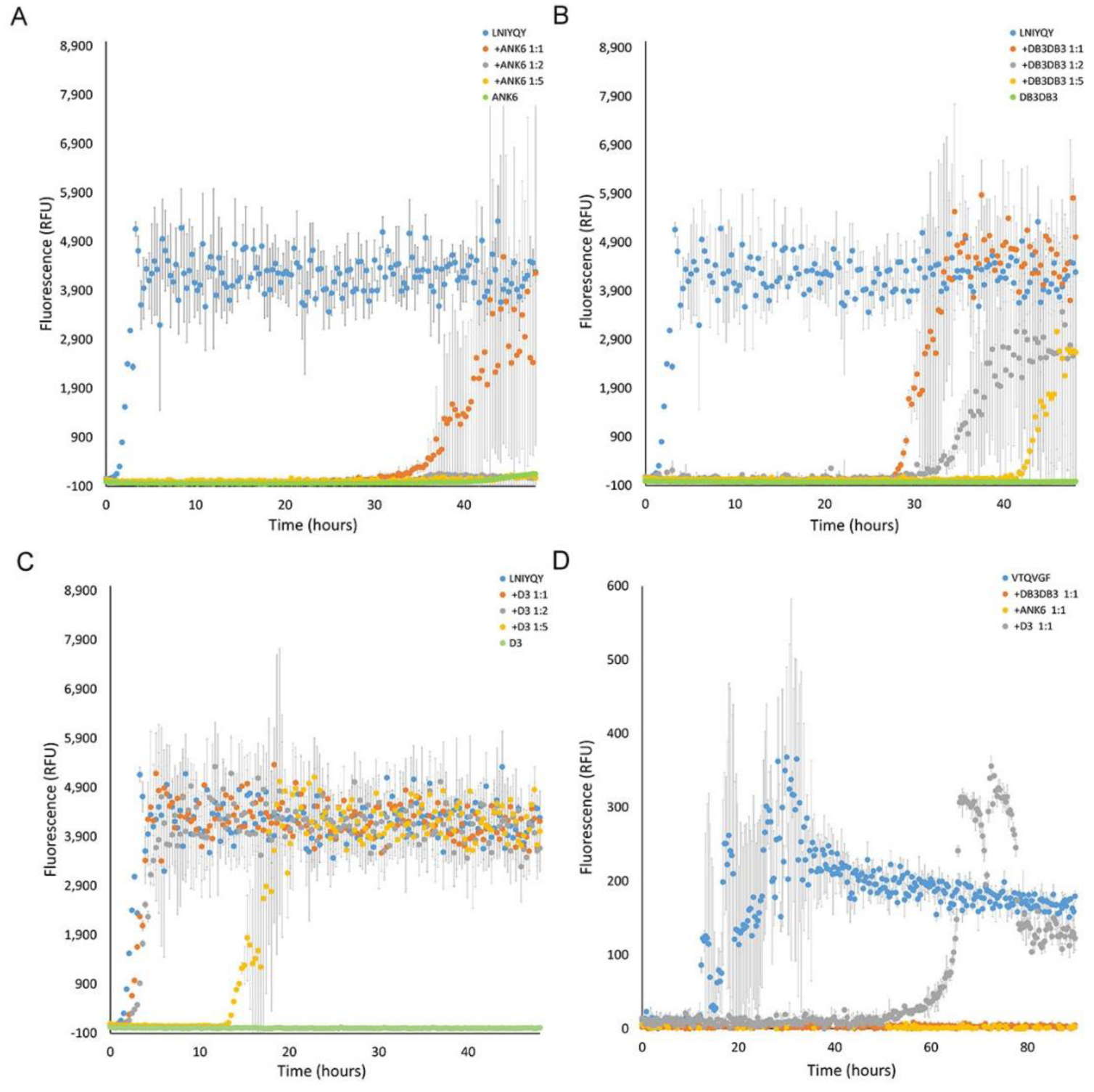
D-peptides inhibit fibrillation of CsgA spine segments. The graphs show mean fluorescence readings of triplicate ThT measurements of CsgA spine segments in the presence of the D-peptide inhibitors. Fibrillation of 100µM _45_LNIYQY_50_ was assessed with 0, 100, 200 and 500 µM ANK6 (**A**), DB3DB3 (**B**) or D3 (**C**) peptides. (**D**) Fibrillation of 500µM _137_VTQVGF_142_ at was examined with 500µM of ANK6, DB3DB3 or D3 peptides. Error bars represent standard error of the mean calculated from triplicates. Each graph represents at least three independent experiments.

**Figure S11.**
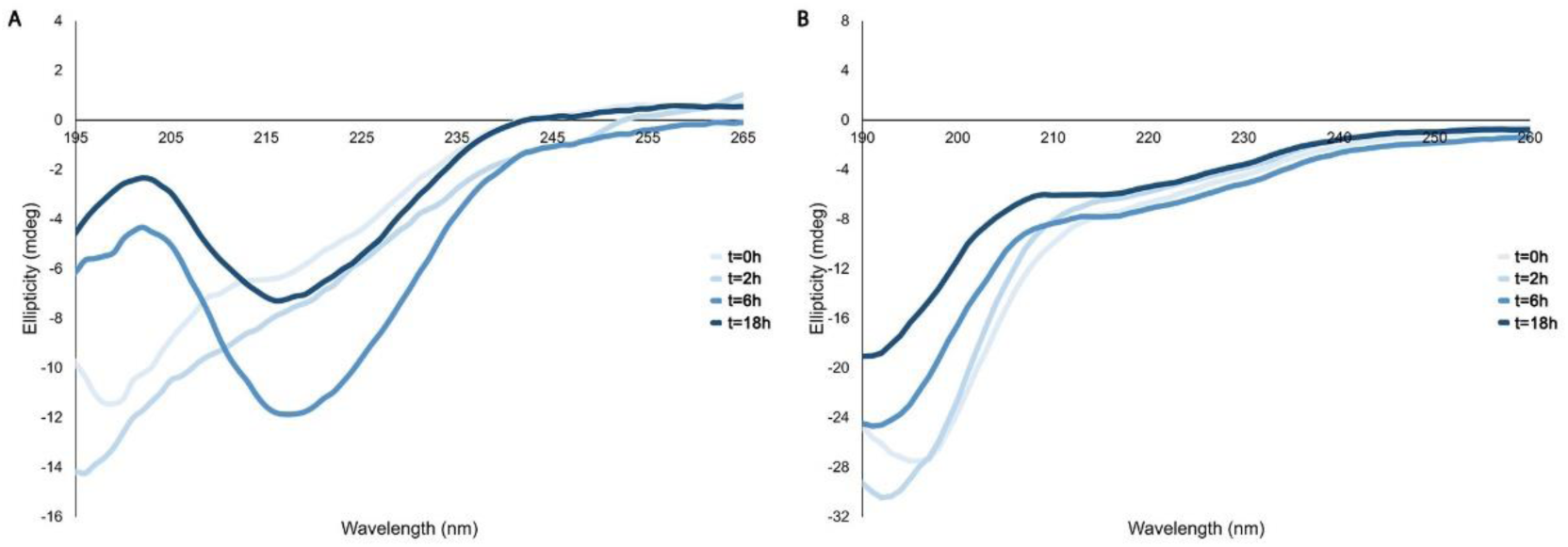
ANK6 prevents secondary structure transitions of CsgA. Time-dependent CD spectra of CsgA incubated alone (**A**) or in the presence of ANK6 (**B**) (1:5 molar ratio). The changes in ellipticity are shown along a wavelength range of 190-265 nm.

**Figure S12.**
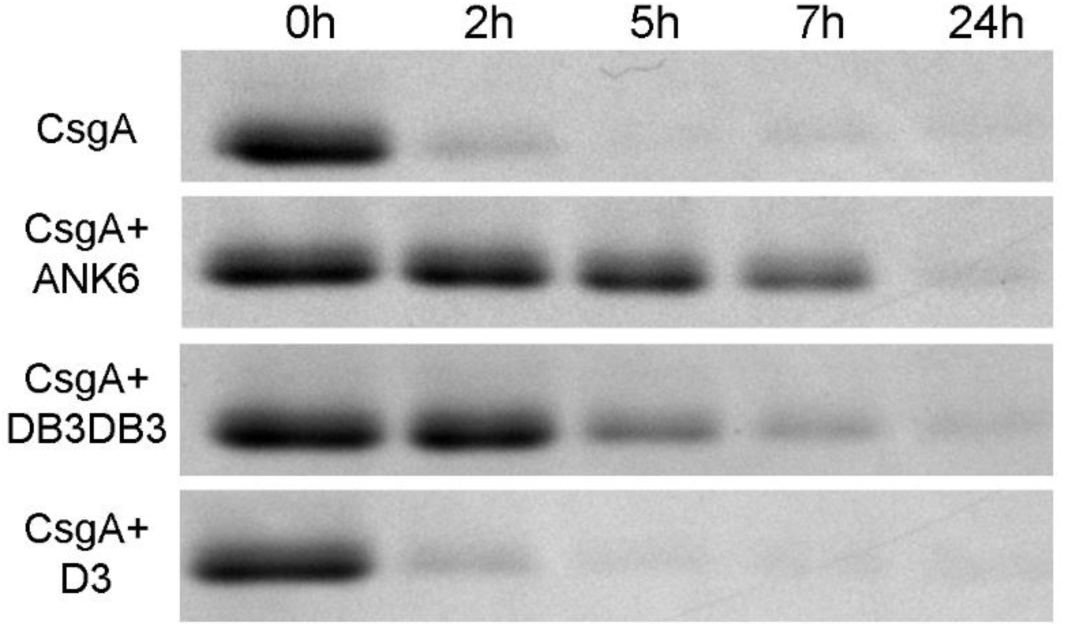
The D-peptides prolong soluble state of CsgA. Migration of soulble CsgA detected by Coomassie-blue staining of 15% SDS-PAGE gel. Incubated CsgA forms insoluble fibrils and does not migrate on the gel compared to freshly purified CsgA. CsgA incubated with ANK6 and DB3DB3 at 1:5 molar ratios showed a prolonged soluble state of CsgA, indicating inhibition of the formation of insoluble fibrils.

**Figure S13.**
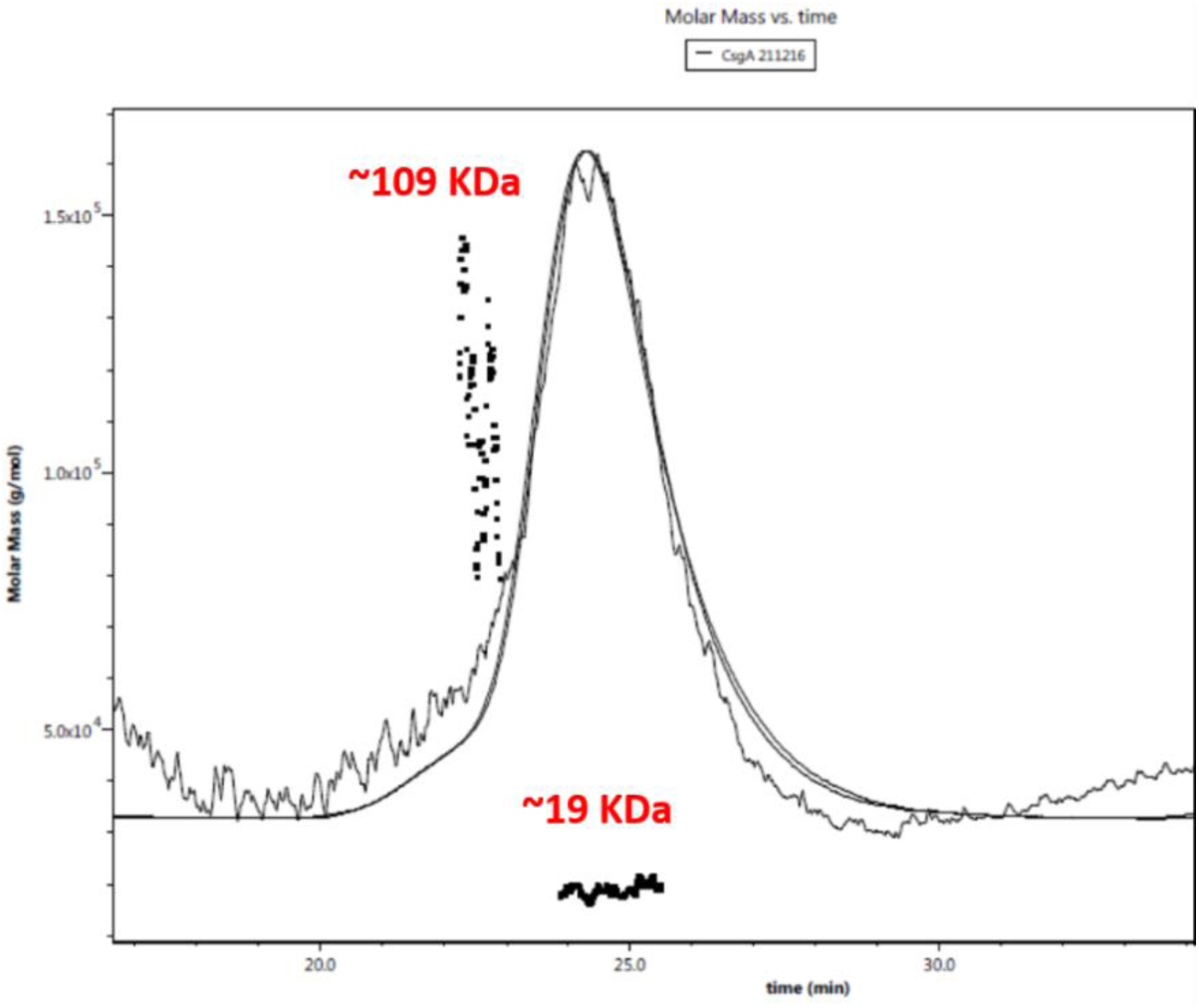
SEC-MALS analysis of freshly purified CsgA. SEC-MALS chromatogram of CsgA presents two main populations with different molecular weights. The major peak corresponds to monomeric CsgA, while the minor peak corresponds to hexamers of CsgA.

**Figure S14.**
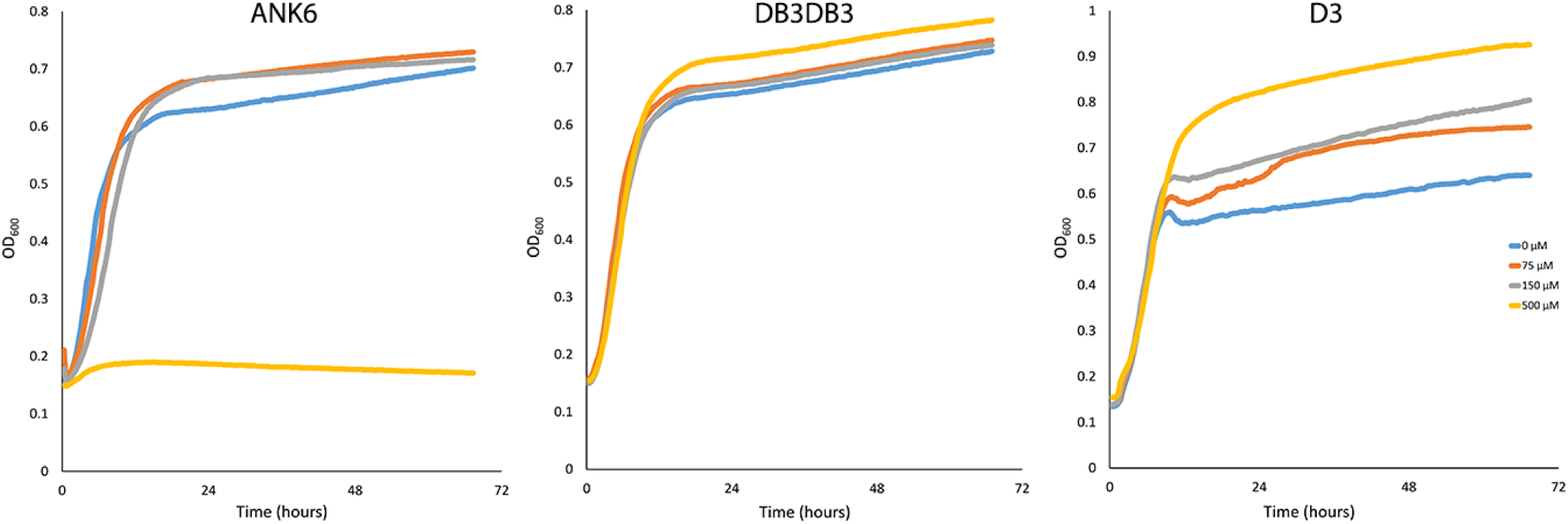
Effect of D-peptides on the growth of *S. typhimurium*. Bacterial growth of the MAE52 strain was assessed in the presence of the D-peptides at concentrations of 0, 75, 150 and 500 µM by a spectrophotometer at OD_600_. An average of triplicate readings were recorded every 20 minutes. No significant effect on the growth phase was evident except with 500 µM of ANK6.

**Figure S15.**
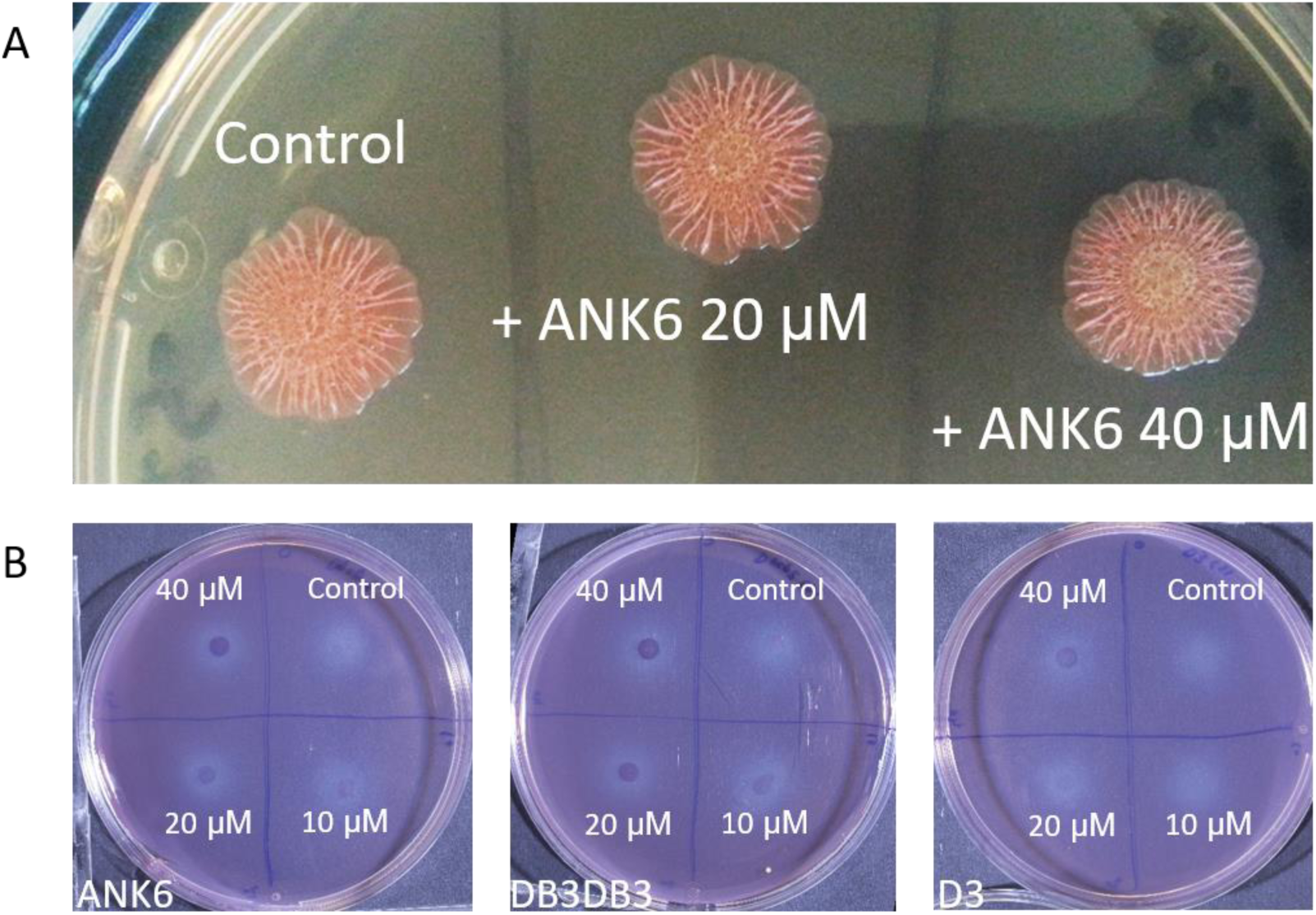
Effect of D-peptides on *S. typhimurium* biofilm grown on CR-supplemented agar. (**A**) *S. typhimurium* MAE52 strain grown on plates with CR-supplemented agar show reddish biofilm colony that adsorbed the dye (left colony on the image). The addition of ANK6 at 20 µM (middle colony) or 40 µM (right colony) show a dose-dependent discoloration at the center of the colony where the drop of the bacteria and D-peptide suspension was placed, indicating less CR adsorption. (**B**) *S. typhimurium* MAE150 strain (cellulose deficient mutant) colonies were grown on plates with CR-supplemented agar; the images depict residual stain on the agar following the removal of the biofilm colonies. From left to right: ANK6, DB3DB3 and D3 added at different concentrations in a clockwise manner, starting in the upper-right corner (0, 10, 20 and 40 µM). Each plate represents a triplicate of repeats and the phenomenon was exhibited in three independent experiments. Quantification of the residual stain is shown in Fig. 6.

**Table S1.**
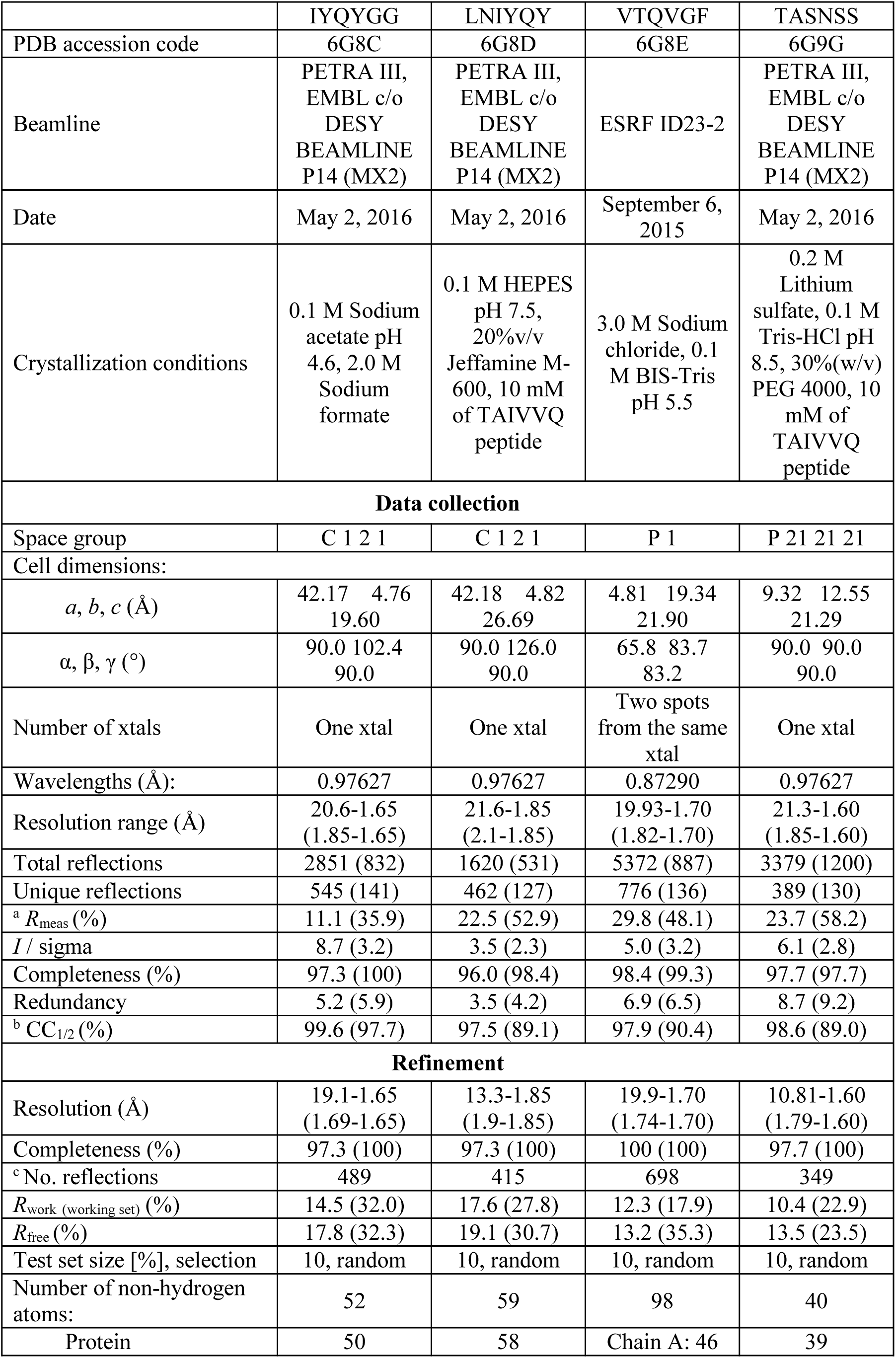
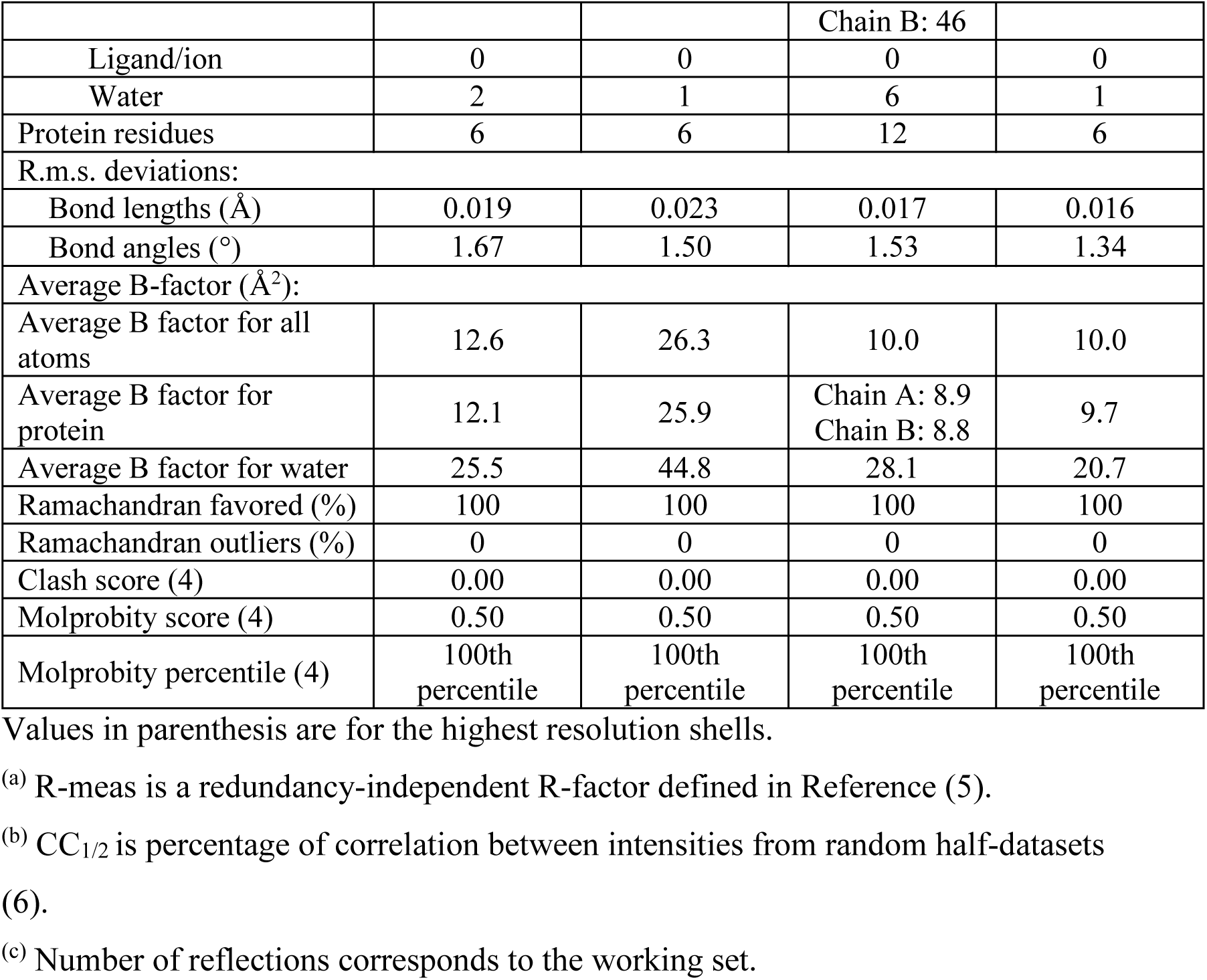
Data collection and refinement statistics (molecular replacement)

**Table S2.**
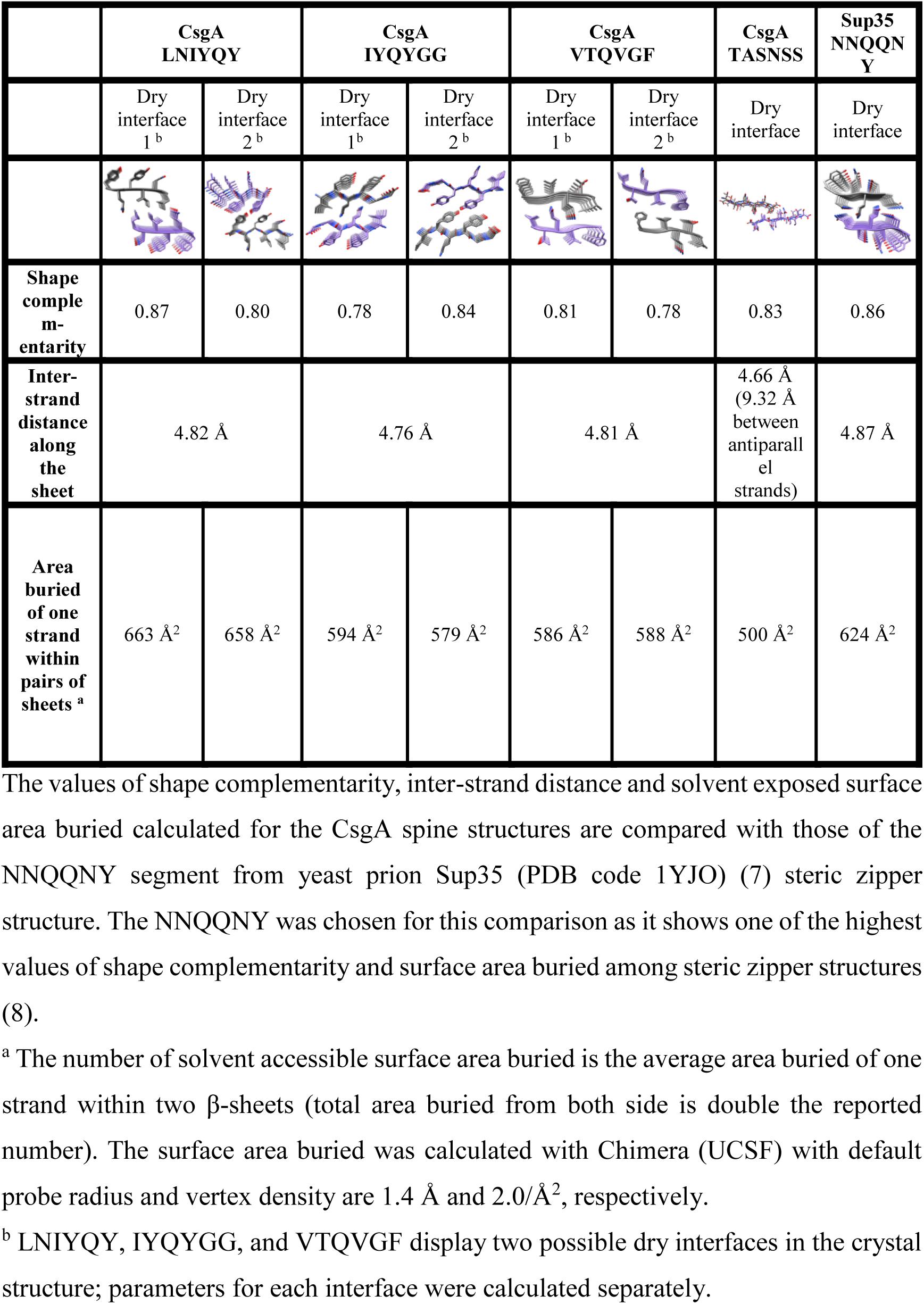
Features of the CsgA spine structures compared to the NNQQNY steric zipper structure.

|  | CsgA<br>LNIYQY |  | CsgA<br>IYQYGG |  | CsgA<br>VTQVGF |  | CsgA<br>TASNSS | Sup35<br>NNQQN<br>Y |
| --- | --- | --- | --- | --- | --- | --- | --- | --- |
|  | Dry<br>interface<br>1<sup>b</sup> | Dry<br>interface<br>2<sup>b</sup> | Dry<br>interface<br>1<sup>b</sup> | Dry<br>interface<br>2<sup>b</sup> | Dry<br>interface<br>1<sup>b</sup> | Dry<br>interface<br>2<sup>b</sup> | Dry<br>interface | Dry<br>interface |
| Shape<br>comple-<br>mentarity | 0.87 | 0.80 | 0.78 | 0.84 | 0.81 | 0.78 | 0.83 | 0.86 |
| Inter-<br>strand<br>distance<br>along<br>the<br>sheet | 4.82 Å |  | 4.76 Å |  | 4.81 Å |  | 4.66 Å<br>(9.32 Å<br>between<br>antiparall<br>el<br>strands) | 4.87 Å |
| Area<br>buried<br>of one<br>strand<br>within<br>pairs of<br>sheets<sup>a</sup> | 663 Å<sup>2</sup> | 658 Å<sup>2</sup> | 594 Å<sup>2</sup> | 579 Å<sup>2</sup> | 586 Å<sup>2</sup> | 588 Å<sup>2</sup> | 500 Å<sup>2</sup> | 624 Å<sup>2</sup> |

The values of shape complementarity, inter-strand distance and solvent exposed surface area buried calculated for the CsgA spine structures are compared with those of the NNQQNY segment from yeast prion Sup35 (PDB code 1YJO) (7) steric zipper structure. The NNQQNY was chosen for this comparison as it shows one of the highest values of shape complementarity and surface area buried among steric zipper structures (8).
<sup>a</sup> The number of solvent accessible surface area buried is the average area buried of one strand within two β-sheets (total area buried from both side is double the reported number). The surface area buried was calculated with Chimera (UCSF) with default probe radius and vertex density are 1.4 Å and 2.0/Å<sup>2</sup>, respectively.
<sup>b</sup> LNIYQY, IYQYGG, and VTQVGF display two possible dry interfaces in the crystal structure; parameters for each interface were calculated separately.

## References

1. Dobson CM (1999) Protein misfolding, evolution and disease. Trends Biochem. Sci. 24(9):329–332.

2. Schmidt M, et al. (2015) Peptide dimer structure in an Abeta(1-42) fibril visualized with cryo-EM. Proc. Natl. Acad. Sci. U.S.A. 112(38):11858–11863.

3. Walti MA, et al. (2016) Atomic-resolution structure of a disease-relevant Abeta(1-42) amyloid fibril. Proc. Natl. Acad. Sci. U.S.A. 113(34):E4976–4984.

4. Colvin MT, et al. (2016) Atomic Resolution Structure of Monomorphic Abeta42 Amyloid Fibrils. J. Am. Chem. Soc. 138(30):9663–9674.

5. Xiao Y, et al. (2015) Abeta(1-42) fibril structure illuminates self-recognition and replication of amyloid in Alzheimer’s disease. Nat. Struct. Mol. Biol. 22(6):499–505.

6. Paravastu AK, Leapman RD, Yau WM, & Tycko R (2008) Molecular structural basis for polymorphism in Alzheimer’s β-amyloid fibrils. Proc. Natl. Acad. Sci. U. S. A. 105(47):18349–18354.

7. Petkova AT, Yau WM, & Tycko R (2006) Experimental constraints on quaternary structure in Alzheimer’s beta-amyloid fibrils. Biochemistry 45(2):498–512.

8. Gremer L, et al. (2017) Fibril structure of amyloid-ss(1-42) by cryoelectron microscopy. Science.

9. Fitzpatrick AWP, et al. (2017) Cryo-EM structures of tau filaments from Alzheimer’s disease. Nature 547(7662):185–190.

10. Van Melckebeke H, et al. (2010) Atomic-resolution three-dimensional structure of HET-s(218-289) amyloid fibrils by solid-state NMR spectroscopy. J. Am. Chem. Soc. 132(39):13765–13775.

11. Chen SW, et al. (2015) Structural characterization of toxic oligomers that are kinetically trapped during alpha-synuclein fibril formation. Proc. Natl. Acad. Sci. U.S.A. 112(16):E1994–2003.

12. Sawaya MR, et al. (2007) Atomic structures of amyloid cross-beta spines reveal varied steric zippers. Nature 447(7143):453–457.

13. Nelson R, et al. (2005) Structure of the cross-beta spine of amyloid-like fibrils. Nature 435(7043):773–778.

14. Wiltzius JJW, et al. (2009) Molecular mechanisms for protein-encoded inheritance. Nat. Struct. Mol. Biol. 16(9):973–U998.

15. Colletier J-P, et al. (2011) Molecular basis for amyloid-beta polymorphism. Proc. Natl. Acad. Sci. U.S.A. 108(41):16938–16943.

16. Rodriguez JA, et al. (2015) Structure of the toxic core of alpha-synuclein from invisible crystals. Nature 525(7570):486–490.

17. Landau M, et al. (2011) Towards a pharmacophore for amyloid. PLoS Biol. 9(6):e1001080.

18. Laganowsky A, et al. (2012) Atomic view of a toxic amyloid small oligomer. Science 335(6073):1228–1231.

19. Laganowsky A, et al. (2010) Crystal structures of truncated alphaA and alphaB crystallins reveal structural mechanisms of polydispersity important for eye lens function. Protein Sci. 19(5):1031–1043.

20. Brumshtein B, et al. (2014) Formation of amyloid fibers by monomeric light chain variable domains. J. Biol. Chem. 289(40):27513–27525.

21. Eisenberg DS & Sawaya MR (2017) Structural Studies of Amyloid Proteins at the Molecular Level. Annu. Rev. Biochem. 86:69–95.

22. Chapman MR, et al. (2002) Role of Escherichia coli curli operons in directing amyloid fiber formation. Science 295(5556):851–855.

23. DePas WH & Chapman MR (2012) Microbial manipulation of the amyloid fold. Res. Microbiol. 163(9–10):592–606.

24. Barnhart MM & Chapman MR (2006) Curli biogenesis and function. Annu. Rev. Microbiol. 60:131–147.

25. Reichhardt C, et al. (2015) Congo Red Interactions with Curli-Producing E. coli and Native Curli Amyloid Fibers. PloS one 10(10):e0140388.

26. Blanco LP, Evans ML, Smith DR, Badtke MP, & Chapman MR (2012) Diversity, biogenesis and function of microbial amyloids. Trends. Microbiol. 20(2):66–73.

27. Taylor JD & Matthews SJ (2015) New insight into the molecular control of bacterial functional amyloids. Front. Cell Infect. Microbiol. 5:33.

28. Evans ML, et al. (2015) The bacterial curli system possesses a potent and selective inhibitor of amyloid formation. Mol. Cell 57(3):445–455.

29. Hammar M, Bian Z, & Normark S (1996) Nucleator-dependent intercellular assembly of adhesive curli organelles in Escherichia coli. Proc. Natl. Acad. Sci. U.S.A. 93(13):6562–6566.

30. Schwartz K & Boles BR (2013) Microbial amyloids–functions and interactions within the host. Curr. Opin. Microbiol. 16(1):93–99.

31. Syed AK & Boles BR (2014) Fold modulating function: bacterial toxins to functional amyloids. Front. Microbiol. 5:401.

32. Soragni A, et al. (2015) Toxicity of eosinophil MBP is repressed by intracellular crystallization and promoted by extracellular aggregation. Mol. Cell 57(6):1011–1021.

33. Maji SK, et al. (2009) Functional amyloids as natural storage of peptide hormones in pituitary secretory granules. Science 325(5938):328–332.

34. Lo AW, Moonens K, & Remaut H (2013) Chemical attenuation of pilus function and assembly in Gram-negative bacteria. Curr. Opin. Microbiol. 16(1):85–92.

35. Tayeb-Fligelman E, et al. (2017) The cytotoxic Staphylococcus aureus PSMalpha3 reveals a cross-alpha amyloid-like fibril. Science 355(6327):831–833.

36. Salinas N, Colletier JP, Moshe A, & Landau M (2018) Extreme amyloid polymorphism in Staphylococcus aureus virulent PSMalpha peptides. Nat. Commun. 9(1):3512.

37. Schwartz K, Syed AK, Stephenson RE, Rickard AH, & Boles BR (2012) Functional amyloids composed of phenol soluble modulins stabilize Staphylococcus aureus biofilms. PLoS Pathog. 8(6):e1002744.

38. Landau M (2018) Mimicking cross-alpha amyloids. Nat. Chem. Biol. 14(9):833–834.

39. Deshmukh M, Evans ML, & Chapman MR (2018) Amyloid by Design: Intrinsic Regulation of Microbial Amyloid Assembly. J. Mol. Biol.

40. Collinson SK, Emody L, Muller KH, Trust TJ, & Kay WW (1991) Purification and characterization of thin, aggregative fimbriae from Salmonella enteritidis. J. Bacteriol. 173(15):4773–4781.

41. Collinson SK, Parker JM, Hodges RS, & Kay WW (1999) Structural predictions of AgfA, the insoluble fimbrial subunit of Salmonella thin aggregative fimbriae. J. Mol. Biol. 290(3):741–756.

42. Shewmaker F, et al. (2009) The functional curli amyloid is not based on in-register parallel beta-sheet structure. J. Biol. Chem. 284(37):25065–25076.

43. Wang X & Chapman MR (2008) Sequence determinants of bacterial amyloid formation. J. Mol. Biol. 380(3):570–580.

44. Wang X, Zhou Y, Ren JJ, Hammer ND, & Chapman MR (2010) Gatekeeper residues in the major curlin subunit modulate bacterial amyloid fiber biogenesis. Proc. Natl. Acad. Sci. U.S.A. 107(1):163–168.

45. Wang X, Smith DR, Jones JW, & Chapman MR (2007) In vitro polymerization of a functional Escherichia coli amyloid protein. J. Biol. Chem. 282(6):3713–3719.

46. Rogers DR (1965) Screening for the amyloid with the thioflavin-T fluorescent method. Am. J. Clin. Pathol. 44:59–61.

47. Dueholm MS, et al. (2011) Fibrillation of the major curli subunit CsgA under a wide range of conditions implies a robust design of aggregation. Biochemistry 50(39):8281–8290.

48. Schubeis T, et al. (2015) Untangling a Repetitive Amyloid Sequence: Correlating Biofilm-Derived and Segmentally Labeled Curli Fimbriae by Solid-State NMR Spectroscopy. Angew. Chem. Int. Ed. Engl. 54(49):14669–14672.

49. Shu Q, et al. (2016) Solution NMR structure of CsgE: Structural insights into a chaperone and regulator protein important for functional amyloid formation. Proc. Natl. Acad. Sci. U.S.A. 113(26):7130–7135.

50. Schubeis T, et al. (2018) Structural and functional characterization of the Curli adaptor protein CsgF. FEBS Lett. 592(6):1020–1029.

51. Taylor JD, et al. (2011) Atomic resolution insights into curli fiber biogenesis. Structure 19(9):1307–1316.

52. Taylor JD, et al. (2016) Electrostatically-guided inhibition of Curli amyloid nucleation by the CsgC-like family of chaperones. Sci. Rep. 6:24656.

53. Goyal P, et al. (2014) Structural and mechanistic insights into the bacterial amyloid secretion channel CsgG. Nature 516(7530):250–253.

54. Cao B, et al. (2014) Structure of the nonameric bacterial amyloid secretion channel. Proc. Natl. Acad. Sci. U.S.A. 111(50):E5439–5444.

55. Leithold LH, et al. (2016) Pharmacokinetic Properties of a Novel D-Peptide Developed to be Therapeutically Active Against Toxic beta-Amyloid Oligomers. Pharm Res 33(2):328–336.

56. van Groen T, Kadish I, Funke SA, Bartnik D, & Willbold D (2013) Treatment with D3 removes amyloid deposits, reduces inflammation, and improves cognition in aged AbetaPP/PS1 double transgenic mice. J. Alzheimers Dis. 34(3):609–620.

57. van Groen T, Kadish I, Funke A, Bartnik D, & Willbold D (2012) Treatment with Abeta42 binding D-amino acid peptides reduce amyloid deposition and inflammation in APP/PS1 double transgenic mice. Adv. Protein Chem. Struct. Biol. 88:133–152.

58. Aileen Funke S, et al. (2010) Oral treatment with the d-enantiomeric peptide D3 improves the pathology and behavior of Alzheimer’s Disease transgenic mice. ACS Chem. Neurosci. 1(9):639–648.

59. van Groen T, et al. (2008) Reduction of Alzheimer’s disease amyloid plaque load in transgenic mice by D3, A D-enantiomeric peptide identified by mirror image phage display. ChemMedChem 3(12):1848–1852.

60. Klein AN, et al. (2017) Optimization of d-Peptides for Abeta Monomer Binding Specificity Enhances Their Potential to Eliminate Toxic Abeta Oligomers. ACS Chem. Neurosci. 8(9):1889–1900.

61. Elfgen A, Santiago-Schubel B, Gremer L, Kutzsche J, & Willbold D (2017) Surprisingly high stability of the Abeta oligomer eliminating all-d-enantiomeric peptide D3 in media simulating the route of orally administered drugs. Eur J Pharm Sci 107:203–207.

62. Jiang N, et al. (2016) Blood-brain barrier penetration of an Abeta-targeted, arginine-rich, d-enantiomeric peptide. Biochim. Biophys. Acta 1858(11):2717–2724.

63. Klein AN, et al. (2016) Optimization of the All-D Peptide D3 for Abeta Oligomer Elimination. PloS one 11(4):e0153035.

64. Sun N, Funke SA, & Willbold D (2012) A survey of peptides with effective therapeutic potential in Alzheimer’s disease rodent models or in human clinical studies. Mini Rev Med Chem 12(5):388–398.

65. Wiesehan K, et al. (2008) Inhibition of cytotoxicity and amyloid fibril formation by a D-amino acid peptide that specifically binds to Alzheimer’s disease amyloid peptide. Protein Eng Des Sel 21(4):241–246.

66. Goldschmidt L, Teng PK, Riek R, & Eisenberg D (2010) Identifying the amylome, proteins capable of forming amyloid-like fibrils. Proc. Natl. Acad. Sci. U.S.A. 107(8):3487–3492.

67. Tartaglia GG, et al. (2008) Prediction of aggregation-prone regions in structured proteins. J. Mol. Biol. 380(2):425–436.

68. Fernandez-Escamilla AM, Rousseau F, Schymkowitz J, & Serrano L (2004) Prediction of sequence-dependent and mutational effects on the aggregation of peptides and proteins. Nat. Biotechnol. 22(10):1302–1306.

69. Maurer-Stroh S, et al. (2010) Exploring the sequence determinants of amyloid structure using position-specific scoring matrices. Nat. Methods 7(3):237–242.

70. Rousseau F, Schymkowitz J, & Serrano L (2006) Protein aggregation and amyloidosis: confusion of the kinds? Curr Opin Struct Biol 16(1):118–126.

71. Sarroukh R, Goormaghtigh E, Ruysschaert JM, & Raussens V (2013) ATR-FTIR: a "rejuvenated" tool to investigate amyloid proteins. Biochim. Biophys. Acta 1828(10):2328–2338.

72. Zandomeneghi G, Krebs MR, McCammon MG, & Fandrich M (2004) FTIR reveals structural differences between native beta-sheet proteins and amyloid fibrils. Protein Sci. 13(12):3314–3321.

73. Moran SD & Zanni MT (2014) How to Get Insight into Amyloid Structure and Formation from Infrared Spectroscopy. The Journal of Physical Chemistry Letters 5(11):1984–1993.

74. Hartman K, et al. (2013) Bacterial curli protein promotes the conversion of PAP248-286 into the amyloid SEVI: cross-seeding of dissimilar amyloid sequences. PeerJ 1:e5.

75. Wasmer C, et al. (2010) Structural similarity between the prion domain of HET-s and a homologue can explain amyloid cross-seeding in spite of limited sequence identity. J. Mol. Biol. 402(2):311–325.

76. Hammer ND, Schmidt JC, & Chapman MR (2007) The curli nucleator protein, CsgB, contains an amyloidogenic domain that directs CsgA polymerization. Proc. Natl. Acad. Sci. U.S.A. 104(30):12494–12499.

77. Andersson EK, et al. (2013) Modulation of Curli Assembly and Pellicle Biofilm Formation by Chemical and Protein Chaperones. Chem Biol 20(10):1245–1254.

78. Wolff M, et al. (2017) Abeta42 pentamers/hexamers are the smallest detectable oligomers in solution. Sci. Rep. 7(1):2493.

79. Reichhardt C, et al. (2016) Influence of the amyloid dye Congo red on curli, cellulose, and the extracellular matrix in E. coli during growth and matrix purification. Anal Bioanal Chem 408(27):7709–7717.

80. Nelson R & Eisenberg D (2006) Recent atomic models of amyloid fibril structure. Curr. Opin. Struc. Biol. 16(2):260–265.

81. Eisenberg D & Jucker M (2012) The amyloid state of proteins in human diseases. Cell 148(6):1188–1203.

82. Jahn TR, et al. (2010) The common architecture of cross-beta amyloid. J. Mol. Biol. 395(4):717–727.

83. Knowles TP, et al. (2007) Role of Intermolecular Forces in Defining Material Properties of Protein Nanofibrils. Science 318(5858):1900.

84. Tian P, et al. (2015) Structure of a Functional Amyloid Protein Subunit Computed Using Sequence Variation. J. Am. Chem. Soc. 137(1):22–25.

85. Cerf E, et al. (2009) Antiparallel beta-sheet: a signature structure of the oligomeric amyloid beta-peptide. Biochem. J. 421(3):415–423.

86. Sarroukh R, et al. (2011) Transformation of amyloid beta(1-40) oligomers into fibrils is characterized by a major change in secondary structure. Cell. Mol. Life Sci. 68(8):1429–1438.

87. Celej MS, et al. (2012) Toxic prefibrillar alpha-synuclein amyloid oligomers adopt a distinctive antiparallel beta-sheet structure. Biochem. J. 443(3):719–726.

88. Gustot A, et al. (2013) Activation of innate immunity by lysozyme fibrils is critically dependent on cross-beta sheet structure. Cell. Mol. Life Sci. 70(16):2999–3012.

89. Richardson JS (1981) The anatomy and taxonomy of protein structure. Adv. Protein Chem. 34:167–339.

90. Wasmer C, et al. (2008) Amyloid fibrils of the HET-s(218-289) prion form a beta solenoid with a triangular hydrophobic core. Science 319(5869):1523–1526.

91. Berthelot K, Lecomte S, Gean J, Immel F, & Cullin C (2010) A yeast toxic mutant of HET-s((218-289)) prion displays alternative intermediates of amyloidogenesis. Biophys. J. 99(4):1239–1246.

92. Fändrich M, et al. (2003) Myoglobin forms amyloid fibrils by association of unfolded polypeptide segments. Proc. Natl. Acad. Sci. U. S. A. 100(26):15463–15468.

93. Moran SD, et al. (2012) Two-dimensional IR spectroscopy and segmental (13)C labeling reveals the domain structure of human γD-crystallin amyloid fibrils. Proc. Natl. Acad. Sci. U.S.A. 109(9):3329–3334.

94. Pinkner JS, et al. (2006) Rationally designed small compounds inhibit pilus biogenesis in uropathogenic bacteria. Proc. Natl. Acad. Sci. U.S.A. 103(47):17897–17902.

95. Cegelski L, et al. (2009) Small-molecule inhibitors target Escherichia coli amyloid biogenesis and biofilm formation. Nat. Chem. Biol. 5(12):913–919.

96. Payne DE, et al. (2013) Tannic acid inhibits Staphylococcus aureus surface colonization in an IsaA-dependent manner. Infect. Immun. 81(2):496–504.

97. Chorell E, et al. (2012) Design and synthesis of fluorescent pilicides and curlicides: bioactive tools to study bacterial virulence mechanisms. Chemistry 18(15):4522–4532.

98. Horvath I, et al. (2012) Mechanisms of protein oligomerization: inhibitor of functional amyloids templates alpha-synuclein fibrillation. J. Am. Chem. Soc. 134(7):3439–3444.

99. Cherny I, et al. (2005) The Formation of Escherichia coli Curli Amyloid Fibrils is Mediated by Prion-like Peptide Repeats. J. Mol. Biol. 352(2):245–252.

100. Friedland RP (2015) Mechanisms of molecular mimicry involving the microbiota in neurodegeneration. J. Alzheimers Dis. 45(2):349–362.

101. Rout SK, Friedmann MP, Riek R, & Greenwald J (2018) A prebiotic template-directed peptide synthesis based on amyloids. Nat. Commun. 9(1):234.

102. Jucker M & Walker LC (2013) Self-propagation of pathogenic protein aggregates in neurodegenerative diseases. Nature 501(7465):45–51.

103. Prusiner SB (1998) Prions. Proc. Natl. Acad. Sci. U. S. A. 95(23):13363–13383.

104. Cui D, Kawano H, Hoshii Y, Liu Y, & Ishihara T (2008) Acceleration of murine AA amyloid deposition by bovine amyloid fibrils and tissue homogenates. Amyloid 15(2):77–83.

105. Miglio A, et al. (2013) Use of milk amyloid A in the diagnosis of subclinical mastitis in dairy ewes. J Dairy Res 80(4):496–502.

106. Villar-Pique A, et al. (2010) Amyloid-like protein inclusions in tobacco transgenic plants. PloS one 5(10):e13625.

107. Pistollato F, et al. (2016) Role of gut microbiota and nutrients in amyloid formation and pathogenesis of Alzheimer disease. Nutr Rev 74(10):624–634.

108. Zhao Y, Dua P, & Lukiw WJ (2015) Microbial Sources of Amyloid and Relevance to Amyloidogenesis and Alzheimer’s Disease (AD). J Alzheimers Dis Parkinsonism 5(1):177.

109. Shoemark DK & Allen SJ (2015) The microbiome and disease: reviewing the links between the oral microbiome, aging, and Alzheimer’s disease. J. Alzheimers Dis. 43(3):725–738.

110. Hill JM & Lukiw WJ (2015) Microbial-generated amyloids and Alzheimer’s disease (AD). Front Aging Neurosci 7:9.

111. Hill JM, Bhattacharjee S, Pogue AI, & Lukiw WJ (2014) The gastrointestinal tract microbiome and potential link to Alzheimer’s disease. Front Neurol 5:43.

112. Sochocka M, et al. (2018) The Gut Microbiome Alterations and Inflammation-Driven Pathogenesis of Alzheimer’s Disease-a Critical Review. Mol Neurobiol.

113. Vogt NM, et al. (2017) Gut microbiome alterations in Alzheimer’s disease. Sci. Rep. 7(1):13537.

114. Harris SA & Harris EA (2015) Herpes Simplex Virus Type 1 and Other Pathogens are Key Causative Factors in Sporadic Alzheimer’s Disease. J. Alzheimers Dis. 48(2):319–353.

115. Zhao Y & Lukiw WJ (2015) Microbiome-generated amyloid and potential impact on amyloidogenesis in Alzheimer’s disease (AD). J Nat Sci 1(7).

116. Miklossy J & McGeer PL (2016) Common mechanisms involved in Alzheimer’s disease and type 2 diabetes: a key role of chronic bacterial infection and inflammation. Aging (Albany NY) 8(4):575–588.

117. Asti A & Gioglio L (2014) Can a bacterial endotoxin be a key factor in the kinetics of amyloid fibril formation? J. Alzheimers Dis. 39(1):169–179.

118. Sharon G, Sampson TR, Geschwind DH, & Mazmanian SK (2016) The Central Nervous System and the Gut Microbiome. Cell 167(4):915–932.

119. Zhou Y, et al. (2012) Promiscuous cross-seeding between bacterial amyloids promotes interspecies biofilms. J. Biol. Chem. 287(42):35092–35103.

120. Lundmark K, Westermark GT, Olsen A, & Westermark P (2005) Protein fibrils in nature can enhance amyloid protein A amyloidosis in mice: Cross-seeding as a disease mechanism. Proc. Natl. Acad. Sci. U.S.A. 102(17):6098–6102.

121. Chen SG, et al. (2016) Exposure to the Functional Bacterial Amyloid Protein Curli Enhances Alpha-Synuclein Aggregation in Aged Fischer 344 Rats and Caenorhabditis elegans. Sci. Rep. 6:34477.

122. Main BS & Minter MR (2017) Microbial Immuno-Communication in Neurodegenerative Diseases. Frontiers in Neuroscience 11:151.

123. Friedland RP & Chapman MR (2017) The role of microbial amyloid in neurodegeneration. PLoS Pathog. 13(12):e1006654.

124. Anthis NJ & Clore GM (2013) Sequence-specific determination of protein and peptide concentrations by absorbance at 205 nm. Protein Sci. 22(6):851–858.

125. Tayeb-Fligelman E & Landau M (2017) X-Ray Structural Study of Amyloid-Like Fibrils of Tau Peptides Bound to Small-Molecule Ligands. Methods Mol. Biol. 1523:89–100.

126. Kabsch W (2010) XDS. Acta Crystallogr. D Biol. Crystallogr. 66(Pt 2):125–132.

127. McCoy AJ, et al. (2007) Phaser crystallographic software. J. Appl. Cryst. 40(4):658–674.

128. Winn MD, et al. (2011) Overview of the CCP4 suite and current developments. Acta Crystallogr. D Biol. Crystallogr. 67(Pt 4):235–242.

129. Murshudov GN, Vagin AA, & Dodson EJ (1997) Refinement of macromolecular structures by the maximum-likelihood method. Acta Crystallogr. D Biol. Crystallogr. 53(Pt 3):240–255.

130. Emsley P, Lohkamp B, Scott WG, & Cowtan K (2010) Features and development of Coot. Acta Crystallogr. D Biol. Crystallogr. 66(Pt 4):486–501.

131. Pettersen EF, et al. (2004) UCSF Chimera–a visualization system for exploratory research and analysis. J. Comput. Chem. 25(13):1605–1612.

132. Lawrence MC & Colman PM (1993) Shape complementarity at protein/protein interfaces. J. Mol. Biol. 234(4):946–950.

133. Romling U, Sierralta WD, Eriksson K, & Normark S (1998) Multicellular and aggregative behaviour of Salmonella typhimurium strains is controlled by mutations in the agfD promoter. Mol. Microbiol. 28(2):249–264.

134. Zogaj X, Nimtz M, Rohde M, Bokranz W, & Romling U (2001) The multicellular morphotypes of Salmonella typhimurium and Escherichia coli produce cellulose as the second component of the extracellular matrix. Mol. Microbiol. 39(6):1452–1463.

135. Lidor O, Al-Quntar A, Pesci EC, & Steinberg D (2015) Mechanistic analysis of a synthetic inhibitor of the Pseudomonas aeruginosa LasI quorum-sensing signal synthase. Sci. Rep. 5:16569.

136. Morris CE, Monier J, & Jacques M (1997) Methods for observing microbial biofilms directly on leaf surfaces and recovering them for isolation of culturable microorganisms. Appl Environ Microbiol 63(4):1570–1576.

137. Dastgheyb S, Parvizi J, Shapiro IM, Hickok NJ, & Otto M (2015) Effect of biofilms on recalcitrance of staphylococcal joint infection to antibiotic treatment. J Infect Dis 211(4):641–650.

## Supporting References

1. Sarroukh R, Goormaghtigh E, Ruysschaert JM, & Raussens V (2013) ATR-FTIR: a “rejuvenated” tool to investigate amyloid proteins. Biochim. Biophys. Acta 1828(10):2328–2338.

2. Zandomeneghi G, Krebs MR, McCammon MG, & Fandrich M (2004) FTIR reveals structural differences between native beta-sheet proteins and amyloid fibrils. Protein Sci. 13(12):3314–3321.

3. Moran SD & Zanni MT (2014) How to Get Insight into Amyloid Structure and Formation from Infrared Spectroscopy. The Journal of Physical Chemistry Letters 5(11):1984–1993.

4. Chen VB, et al. (2010) MolProbity: all-atom structure validation for macromolecular crystallography. Acta Crystallogr. D Biol. Crystallogr. 66(Pt 1):12–21.

5. Diederichs K & Karplus PA (1997) Improved R-factors for diffraction data analysis in macromolecular crystallography. Nat. Struct. Biol. 4(4):269–275.

6. Karplus PA & Diederichs K (2012) Linking crystallographic model and data quality. Science 336(6084):1030–1033.

7. Nelson R, et al. (2005) Structure of the cross-beta spine of amyloid-like fibrils. Nature 435(7043):773–778.

8. Sawaya MR, et al. (2007) Atomic structures of amyloid cross-beta spines reveal varied steric zippers. Nature 447(7143):453–457.

